# Loss of CREST leads to neuroinflammatory responses and ALS-like motor defects in mice

**DOI:** 10.1101/260133

**Authors:** Cheng Cheng, Kan Yang, Xinwei Wu, Yuefang Zhang, Shifang Shan, Aaron Gitler, Anirvan Ghosh, Zilong Qiu

## Abstract

Amyotrophic lateral sclerosis (ALS) is a late onset neurodegenerative disease with fast progression. Mutations of the *CREST* gene (also known as *SS18L1*) are identified in sporadic ALS patients. Whether *CREST* mutations may lead to ALS remained largely unclear. In this study, we showed that the ALS-related CREST-Q388X mutation exhibited *loss-of-function* effects. Importantly, we found that microglial activation were prevalent in CREST haploinsufficieny mice and the Q394X mice mimicking the human *CREST* Q388X mutation. Furthermore, we showed that both CREST haploinsufficieny and the Q394X mice displayed deficits in motor coordination. Finally, we identified the critical role of CREST-BRG1 complex in repressing the expression of immune-related cytokines including *Ccl2* and *Cxcl10* in neurons, via histone deacetylation, providing the molecular mechanisms underlying inflammatory responses lack of CREST. These findings indicate that elevated inflammatory responses in a subset of ALS may be caused by neuron-derived factors, suggesting potential therapeutic methods through inflammation pathways.

**In Brief:** Cheng et al. discovered that neuronal loss of CREST reduces the protein level of FUS, de-represses the transcriptional inhibition of chemokine genes which in turn causes microglial activation and proinflammation, and ultimately leads to axonal degeneration of motor neurons and impairment of locomotion.

## INTRODUCTION

Ayotrophic lateral sclerosis (ALS) is one of the most severe neurodegenerative diseases characterized by the fast progressive degeneration of motor neurons in the central nervous system (CNS). Usually within 3–5 years of disease onset, patients suffer multiple symptoms including muscle atrophy, paralysis and respiratory failure, during which there are limited therapeutic approaches to alleviate disease symptoms. Moreover, clinical features of ALS appeared great heterogeneity, meaning that ALS patients may experience motor neuron death onset in different regions of CNS, such as the spinal cord, brainstem or motor cortex (Taylor et al., 2016). ALS has heavy genetic components in which a series of genetic mutations have been identified. Since the ALS-causing mutations in the *SOD1* gene were reported in 1993 (Rosen et al., 1993), more than 50 ALS-associated genes have been subsequently reported, including the *TARDBP*, encoding TAR DNA-binding protein 43 (TDP43) (Sreedharan et al., 2008), *FUS* (Kwiatkowski et al., 2009; Vance et al., 2009), *C9orf72* (DeJesus-Hernandez et al., 2011; Renton et al., 2011). With the development of next-generation sequencing (NGS) technology, many previously unknown ALS-linked mutations have been certified via whole-exome sequencing. Several *de novo* and inherited mutations in *CREST* have been recently reported in ALS patients via NGS-based whole-exome sequencing or target gene sequencing approaches (Chesi et al., 2013; Cirulli et al., 2015; Teyssou et al., 2014), suggesting that *CREST* may be a potential ALS-causing gene. Among all the mutations, we focus on one *de novo* missense mutation, CREST-Q388X, which leads to a truncation that lacks the nine amino acids in the C-terminus (Chesi et al., 2013).

*CREST* is characterized as a calcium-regulated transcriptional activator of which C-terminus is responsible for gene activation through interacting to CREB-binding protein (CBP) (Aizawa et al., 2004). Whereas the N-terminus of CREST protein has auto-inhibitory function via interacting with the chromatin remodeling BRG1 complex which in turn recruits histone deacetylase complex HDAC1 to inhibit gene transcription (Qiu and Ghosh, 2008). Although previous study has shown that both loss of CREST and overexpression of Q394X (the corresponding truncation of mouse *Crest* homolog) or I123M mutant block the depolarization-induced dendritic outgrowth in cultured neurons (Aizawa et al., 2004; Chesi et al., 2013), whether mutations of *CREST* lead to ALS-like phenotypes *in vivo* remain to be determined.

Previous publications have demonstrated that non-neuron-involved chronic inflammatory responses play a critical role in the pathogenesis of ALS (Clement et al., 2003; Glass et al., 2010; Neymotin et al., 2009; Yamanaka et al., 2008). During neuroinflammatory responses, microglial activation, astrogliosis and infiltration of peripheral immune cells are the main observable pathophysiological hallmarks (Alexianu et al., 2001; Beers et al., 2011; Engelhardt and Appel, 1990; Engelhardt et al., 1993; Hall et al., 1998; Kawamata et al., 1992; Mantovani et al., 2009; Turner et al., 2004), and subsequent production of neurotoxic factors such as tumor necrosis factor α (TNF-α) and interleukin 1β (IL-1β) can deteriorate disease progression (Meissner et al., 2010; Weydt et al., 2004). As resident macrophages in CNS, microglia perform the first defense line of innate immune system, and thereby the excess activation of microglia predominantly conducts sustained neuroinflammatory responses that contribute to the progression of ALS or other neurodegenerative diseases (Boillee et al., 2006; Frakes et al., 2014; Mass et al., 2017). Therefore, whether CREST may contribute to the neuroinflammation *in vivo* and mice carrying ALS-like CREST mutations may photocopy ALS-like phenotypes are yet to be determined.

In this study, we show the impaired protein stability of Q388X mutant compared to wild-type (WT) CREST in contrast to the previous report (Kukharsky et al., 2015). Importantly, we demonstrate that microglial appearance exhibits activated morphology including the enlargement of cell bodies and the decreased complexity of processes in both CREST knockout (KO) mice and homozygous *Crest ^Q394X/Q394X^* (Abbreviated as Q394X) mice, suggesting that CREST haploinsufficieny and ALS-related mutation leads to microglial activation and sustains the proinflammatory state in CNS. In agreement with ALS-like phenotypes, we also show the denervation of tibialis anterior (TA) muscles in CREST KO mice and the locomotion impairment in both CREST KO and Q394X mice. Finally, we found the upregulation of two important chemokines, *Cxcl10* and *Ccl2*, in neurons lack of CREST which may contribute to the activation of inflammation in CNS. Taken together, we demonstrate that neuronal loss of CREST function caused by the ALS-linked mutation induces the alteration of immune-related genes expression which leads to the microglial activation and sustained proinflammatory responses, which in turn impair motor neurons and motor behaviors of mutant mice.

## RESULTS

### The Q388X Mutation Decreases Protein Stability of CREST *in vitro* and *in vivo*

To study protein properties of the CREST-Q388X mutant, we constructed lentiviral vectors expressing cDNA of human WT CREST and Q388X mutant respectively, and infected lentivirus into primary cortical neurons isolated from embryonic C57BL/6 mice. Although there was no statistical difference between basal levels of CREST WT and Q388X mutant protein, we surprisingly found that the protein level of Q388X mutant was significantly reduced compared to WT if protein synthesis was blocked by chlorhexidine (CHX) treatment, suggesting that the protein stability of CREST-Q388X mutant is much lower compared to WT in cultured neurons *in vitro* (Figures 1A and 1B).

**Figure 1.**
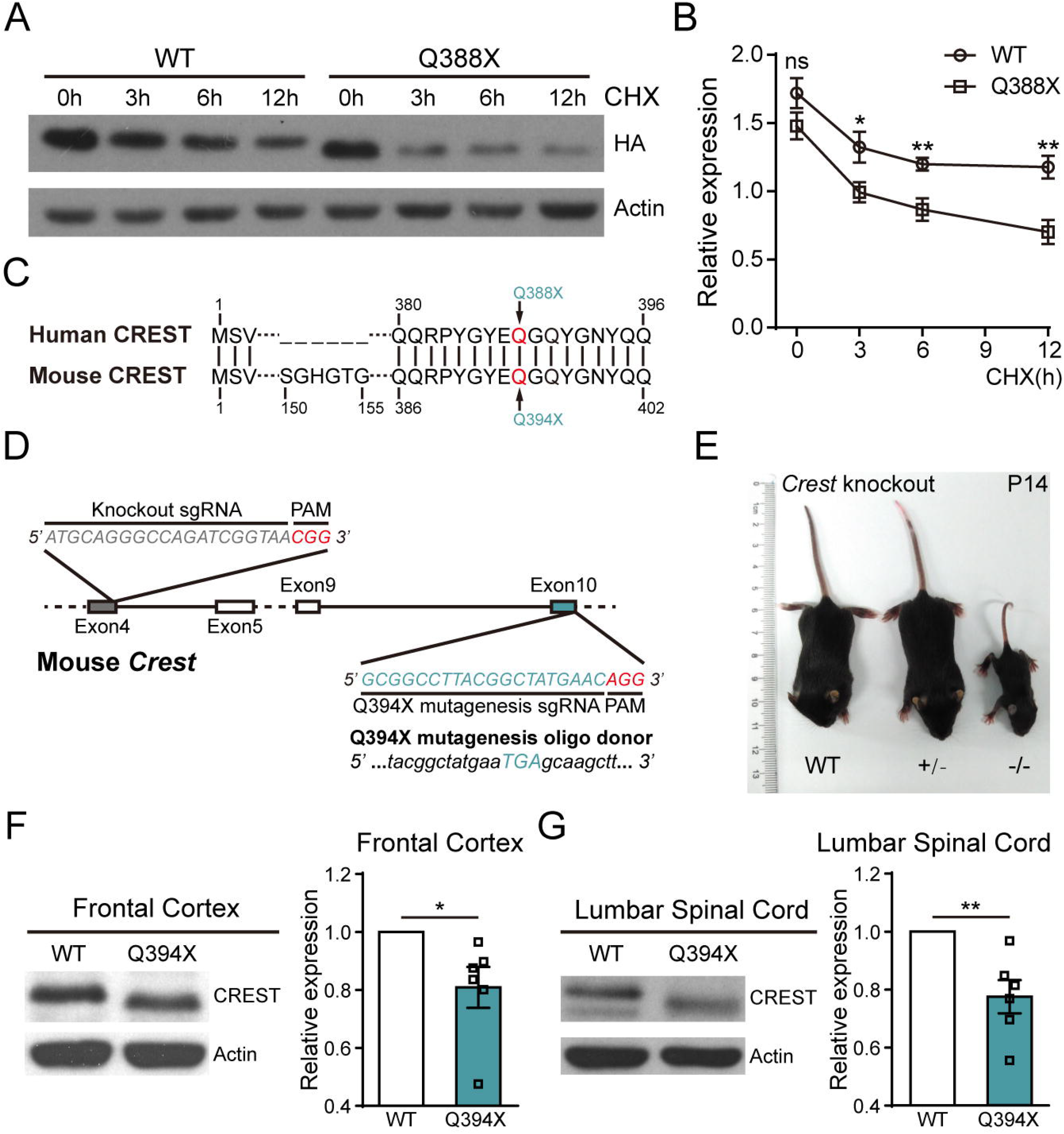
The ALS Mutation Q388X Leads to Instability of CREST Protein *In Vitro* and *In Vivo*. (A and B) Protein levels of WT and Q388X mutant form of CREST in primary cortical neurons infected with lentivirus expressing HA tagged CREST WT or Q388X cDNA respectively, treated with chlorhexidine (CHX, 20μg/ml) for 0h, 3h, 6h, 12h, measured by immunoblot (A) and quantification (B). Relative expression represents the ratios of HA (upper panel) and Actin (lower panel) band intensities determined by Image J. (C) The schematic illustration for partial alignment of human CREST protein with mouse CERST protein, showing the conserved position of ALS mutation (Q388X) and the corresponding site in mouse CREST protein (Q394X). (D) Strategies of the constructing CREST knockout (KO) and CREST Q394X mutation mice using CRISPR/Cas9 technology. (E) Body size of a *Crest ^-/-^* mouse compared to *Crest ^+/-^* and WT littermates at P14. (F and G) Endogenous protein levels of CREST in the frontal cortices (F) and the lumbar spinal cords (G) from 6-month-old Q394X mice and WT littermates (n = 6, each phenotype). Error bars represent SEM. \**p* <0.05, \*\**p* < 0.01, Student’s *t* test.

To further examine the role of CREST Q388X mutation *in vivo*, we constructed CREST knockout (KO) mice and the Q394X mice carrying point mutation Q394X in the *Crest* gene (the corresponding mutation of mouse CREST homolog for human CREST-Q388X) with the CRISPR/Cas9 technology (Figures 1C and 1D). Consistent with previous report, most of the homozygous KO (*Crest ^-/-^*) mice were lethal, and survivors were much smaller in size and had severe locomotion impairment compared to WT or heterozygous (*Crest ^+/-^*) littermates (Figure 1E) (Aizawa et al., 2004). We tested the protein level of endogenous CREST by performing Western blotting on samples collected from frontal cortices and lumbar spinal cords of Q394X and WT mice of 6 months old. Consistently, we found that the protein level of CREST declined about 20% in brain and spinal cord of Q394X mice compared to WT mice, indicating that Q394X mutation leads to instability of endogenous CREST protein *in vivo* (Figures 1F and 1G). Therefore, we suggest that CREST*-*Q388X mutant exhibits *loss-of-function* effects in causing ALS in human patients.

### Activated Morphological Appearance of Microglia and the Denervation of TA Muscles Occur in CREST KO Mice

To establish the causal connection between loss of CREST and ALS pathogenesis, we set out to determine whether loss of CREST caused the ALS-associated pathological features *in vivo*. Since homozygous *Crest ^-/-^* mice could only survive less than one month, we examined histology markers in the brain and lumbar spinal cord sections of *Crest ^-/-^, Crest ^+/-^* and WT mice at postnatal day 14 (P14). First, we found that the number of cholinergic (ChAT-positive) motor neurons in the ventral horn of lumbar spinal cord had no statistical difference between *Crest ^-/-^, Crest ^+/-^* and WT mice (Figures S1A, S1B and S1C). Since inflammatory responses mediated by microglia in the CNS have been implicated in numerous neurodegenerative diseases and motor neuron disorders, we would like to examine whether the neuroinflammation was evoked in the absence of CREST. We measured the inflammatory responses by immunostaining Iba1, the marker for microglia, in cerebral motor cortex and lumbar spinal cord of *Crest ^-/-^, Crest ^+/-^* and WT mice at P14. In terms of the overall cell number of Iba1-positive microglia, we did not detect alteration in either motor cortex or lumbar spinal cord of *Crest ^-/-^*, or *Crest ^+/-^* mice compared to WT littermates (Figures S1D, S1E and S1F). However, we observed that Iba1-positive microglia represented activated morphological features, including enlarged cell bodies and less complex branches, in both lumbar spinal cord and motor cortex of CREST KO mice, compared to WT littermates (Figures S1G-S1P). Moreover, we showed more dramatic decrease of the length and number of microglial branches in lumbar spinal cord and motor cortex of *Crest ^-/-^* mice compared to *Crest ^+/-^* littermates (Figures S1J, S1K, S1O and S1P). It is also worth to mention that there is higher microglial activation level in spinal cord than in motor cortex of *Crest ^+/-^* mice at P14 (Figures S1J, S1K, S1O and S1P).

We further performed immunohistochemistry analysis with Iba1 antibody in aged *Crest ^+/-^* and WT mice at 6 months of age. Consistently, we observed morphological features of chronically activated microglia (increased soma area and decreased branch length or number) in both lumbar spinal cord and motor cortex in *Crest ^+/-^* mice, but not in WT mice of the same age (Figures 2A-2J). Interestingly, we noticed that the complexity of microglial branches did not change in cerebral motor cortex of *Crest ^+/-^* mice at P14, but at 6 months, significant decreases in branch length and number were detected in the same region, suggesting that the extent of microglial activation increases in *Crest ^+/-^* mice with age (Figures 2I, 2J, S1O and S1P). Taken together, we demonstrate that loss of CREST directly leads to microglial activation in the CNS of mice.

**Figure 2.**
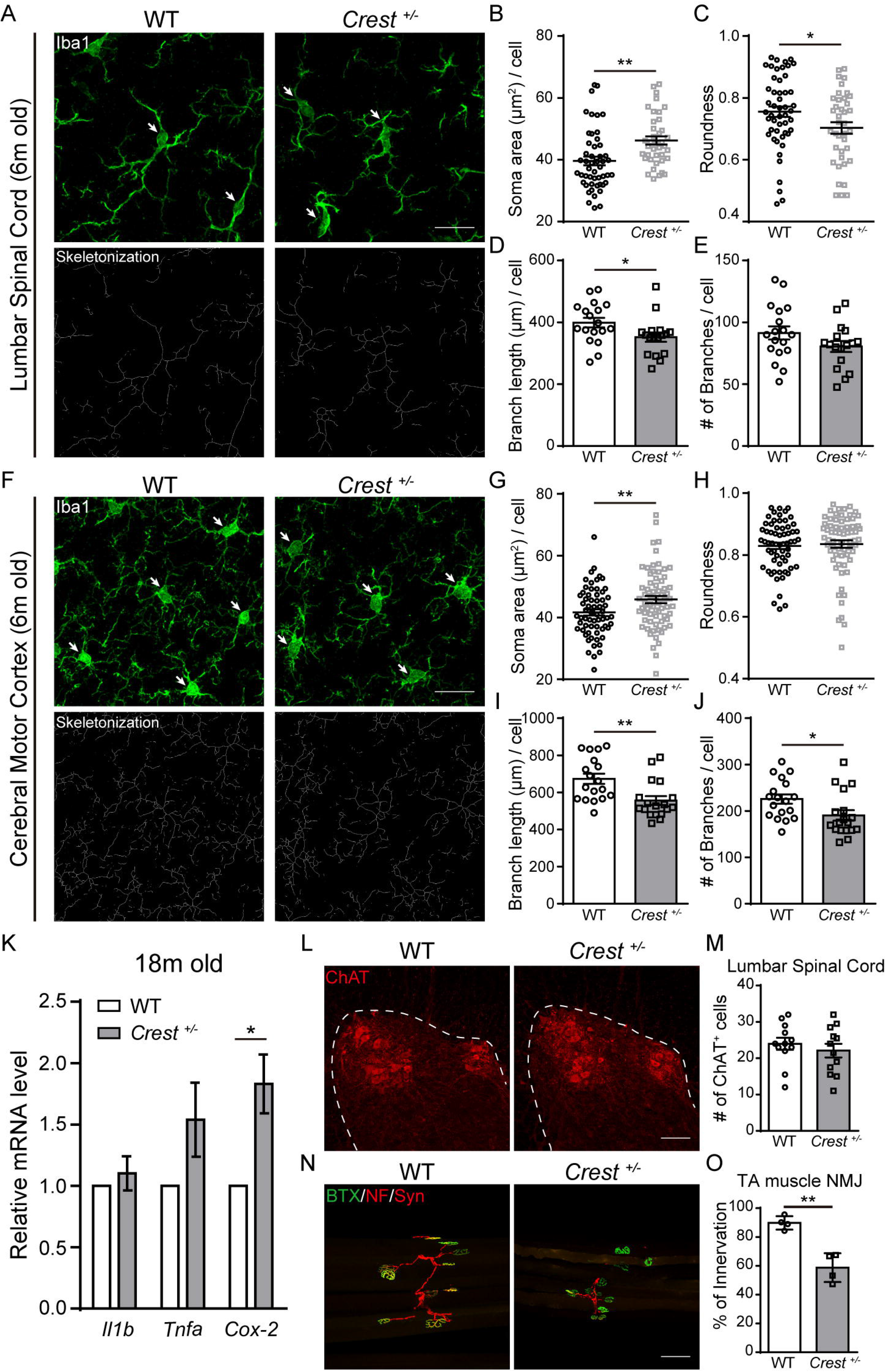
CREST Haploinsufficiency Leads to Upregulation of Inflammatory Responses in the Central Nervous System and the Denervation of Neuromuscular Junctions in TA Muscles. (A) Representative immunohistochemistry images of Iba1-positive microglia (highlighted by arrows, green, upper panels) and their skeletonized appearance (lower panels) in the lumbar spinal cords of 6-month-old *Crest ^+/-^* mice (n=3) and WT littermates (n=3). Scale bar, 20μm. (B-E) Analysis and quantification of microglial morphological parameters including soma area (B) and roundness (C) of the projection of Iba1-positive cell bodies, and branch length (D) and branch number (E) per cell in the lumbar spinal cords of 6-month-old *Crest ^+/-^* mice (n=3) and WT littermates (n=3). (F) Representative immunohistochemistry images of Iba1-positive microglia (highlighted by arrows, green, upper panels) and their skeletonized appearance (lower panels) in the cerebral motor cortices of 6-month-old *Crest ^+/-^* mice (n=3) and WT littermates (n=3). Scale bar, 20μm. (G-J) Analysis and quantification of microglial morphological parameters including soma area (G) and roundness (H) of the projection of Iba1-positive cell bodies, and branch length (I) and branch number (J) per cell in the cerebral motor cortices of 6-month-old *Crest ^+/-^* mice (n=3) and WT littermates (n=3). (K) mRNA levels of proinflammatory genes, such as IL-1β, TNF-α and COX-2, in frontal cortices from 18-month-old *Crest ^+/-^* mice (n=9) and WT littermates (n=3) measured by quantitative RT-PCR. (L and M) Representative immunohistochemistry images (L) and quantification (M) of ChAT-positive (red) motor neurons in ventral horn of the lumbar spinal cords of 6-month-old *Crest ^+/-^* mice (n=3) and WT littermates (n=3). Dashed lines indicate grey matter of lumbar spinal cord. Scale bar, 100μm. (N and O) Representative immunohistochemistry images (N) and quantification (O) of the innervation of neuromuscular junctions in tibialis anterior (TA) muscles of 6-month-old *Crest ^+/-^* mice (n=3) and WT littermates (n=3). Acetylcholine receptors were labeled with α-bungarotoxin (BTX) CF488A conjugate (green), motor neuron axons were labeled with anti-neurofilament-L (NF) antibody (red), and synapses were labeled with anti-synapsin-1 (Syn) antibody (red). Scale bar, 100μm. Error bars represent SEM. \**p* <0.05, \*\**p* < 0.01, Student’s *t* test.

Besides the morphological analysis of microglia, we tested expression levels of genes associated with the neurotoxic proinflammatory state in *Crest ^+/-^* and WT mice. Interestingly, we found a 2-fold upregulation of proinflammatory gene cyclooxygenase-2 (*Cox*-2, also known as *Ptgs2*), but not IL-1β (*Il1b*) or TNF-α (*Tnfa*), in cortical tissue of *Crest* mice at 18 months of age (Figure 2K). This result is consistent with the finding that increased expression of *Cox*-2 in ALS patients and SOD1 mouse model (Almer et al., 2001; McGeer and McGeer, 2002). Together, we demonstrate that loss of CREST results in both microglial activation and the onset of proinflammatory processes in CNS.

We further analyzed neurodegenerative phenotypes of both somatic and axonal morphology of motor neurons in lumbar spinal cord of *Crest ^+/-^* mice and WT littermates at 6 months of age. Although there was no statistical difference in the number of ChAT-positive motor neurons in lumbar ventral horn between WT and *Crest ^+/-^* mice (Figures 2L and 2M), we found that the ratio of innervation of neuromuscular junctions (NMJs) in TA muscles significantly decreased in *Crest ^+/-^* mice compared to WT littermates (Figures 2N and 2O), suggesting the signs of axonal degeneration of motor neurons in lumbar ventral horn of *Crest ^+/-^* mice. Moreover, we intend to determine whether loss of CREST directly leads to impaired axonal morphology in a cell-autonomous manner by examining the effect of CREST in primary cultured neurons *in vitro*. As an important hallmark of ALS, the aberrant accumulation of protein inclusions in cellular cytoplasm may be toxic to neurons (Mackenzie et al., 2007; Neumann et al., 2006; Van Deerlin et al., 2008). However, we did not find that the formation of YB1-positive stress granules under the treatment of sodium arsenite was influenced in isolated primary cortical neurons from embryonic CREST KO or WT littermates (Figures S2A and S2B). To further illustrate the dosage effects of CREST on axonal morphology, we transfected plasmids expressing short hairpin RNA (shRNA) of CREST (Qiu and Ghosh, 2008) or human WT CREST cDNA into mouse primary cultured neurons and detected the axonal phenotype by immunostaining with axonal marker SMI312. We found that neither knockdown nor overexpression of CREST in neurons affected the axonal length (Figures S2C and S2D). Thus, we suggest that the denervation of NMJs in *Crest ^+/-^* mice may be caused by chronic microglial activation and proinflammatory responses, rather than CREST regulating axonal development directly.

### Activation of Microglia Occurs in the CREST Q394X Mice

Next, we would like to determine whether inflammatory responses are also activated in the CNS of Q394X mutant mice. After immunohistochemical analysis of Iba1-positive microglia in cerebral motor cortex and lumbar spinal cord of Q394X mice and WT littermates, we showed significant increase in soma area and decrease in soma roundness but no alteration of branch complexity of microglia in lumbar spinal cord of Q394X mice at 6 months, compared to WT littermates (Figures 3A-3E). Moreover, there were increase of soma area and decrease of length and number of branches of microglia in motor cortex of Q394X mice compared to WT at 6 months (Figures 3F-3J). Therefore, we identified similar signs of increased inflammatory responses in motor cortex and lumbar spinal cord of Q394X mice, as they are in CREST haplosufficiency mice, suggesting that Q394X mice exhibits neuropathological symptoms in CNS.

**Figure 3.**
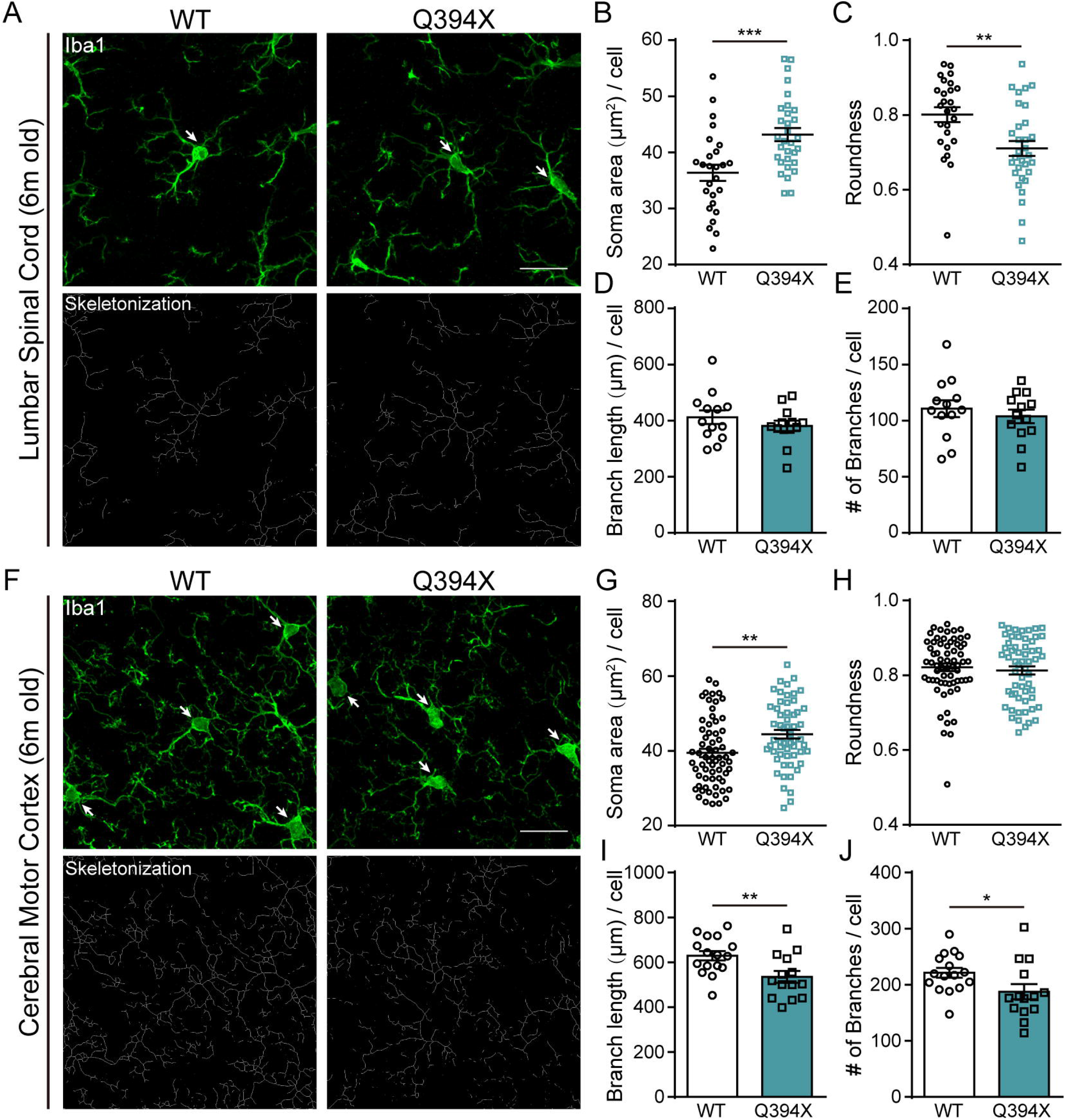
Microglia Are Prevalently Activated in Q394X Mice. (A) Representative immunohistochemistry images of Iba1-positive microglia (highlighted with arrows, green, upper panels) and their skeletonized appearance (lower panels) in the lumbar spinal cords of 6-month-old Q394X mice (n=3) and WT littermates (n=3). Scale bar, 20μm (B-E) Analysis and quantification of microglial morphological parameters including soma area (B) and roundness (C) of the projection of Iba1-positive cell bodies, and branch length (D) and branch number (E) per cell in the lumbar spinal cords of 6-month-old Q394X mice (n=3) and WT littermates (n=3). (F) Representative immunohistochemistry images of Iba1-positive microglia (highlighted with arrows, green, upper panels) and their skeletonized appearance (lower panels) in the cerebral motor cortices of 6-month-old Q394X mice (n=3) and WT littermates (n=3). Scale bar, 20μm (G-J) Analysis and quantification of microglial morphological parameters including soma area (G) and roundness (H) of the projection of Iba1-positive cell bodies, and branch length (I) and branch number (J) per cell in the cerebral motor cortices of 6-month-old Q394X mice (n=3) and WT littermates (n=3). Error bars represent SEM. \**p* <0.05, \*\**p* < 0.01, and \*\*\**p* < 0.001, Student’s *t* test.

### Both CREST KO and Q394X Mice Show the Impairment of Motor Phenotypes

To determine whether *Crest ^+/-^* or Q394X mice may exhibit ALS-like motor defects, we performed a battery of behavioral tests including open field, footprint, grip strength, rotarod and beam walking (Brooks and Dunnett, 2009) on *Crest ^+/-^* mice and WT littermates. *Crest ^+/-^* mice appeared to be generally healthy at 18 months (Figure S3A), and exhibited no defects in open field, footprint and grip strength tests (Figures S3B-S3D). However, we found that *Crest ^+/-^* mice started to exhibit deficits of motor coordination in the accelerating rotarod test from 14 months of age (Figure 4A). Moreover, *Crest ^+/-^* mice fell more easily from the rotating rod than WT littermates in the fixed speed task of rotarod test (Figure 4B). In the beam walking test, *Crest ^+/-^* mice showed significantly more slipped steps compared to WT mice (Figure 4C, Videos S1 and S2), indicating that motor coordination abilities of *Crest ^+/-^* mice are indeed compromised.

**Figure 4.**
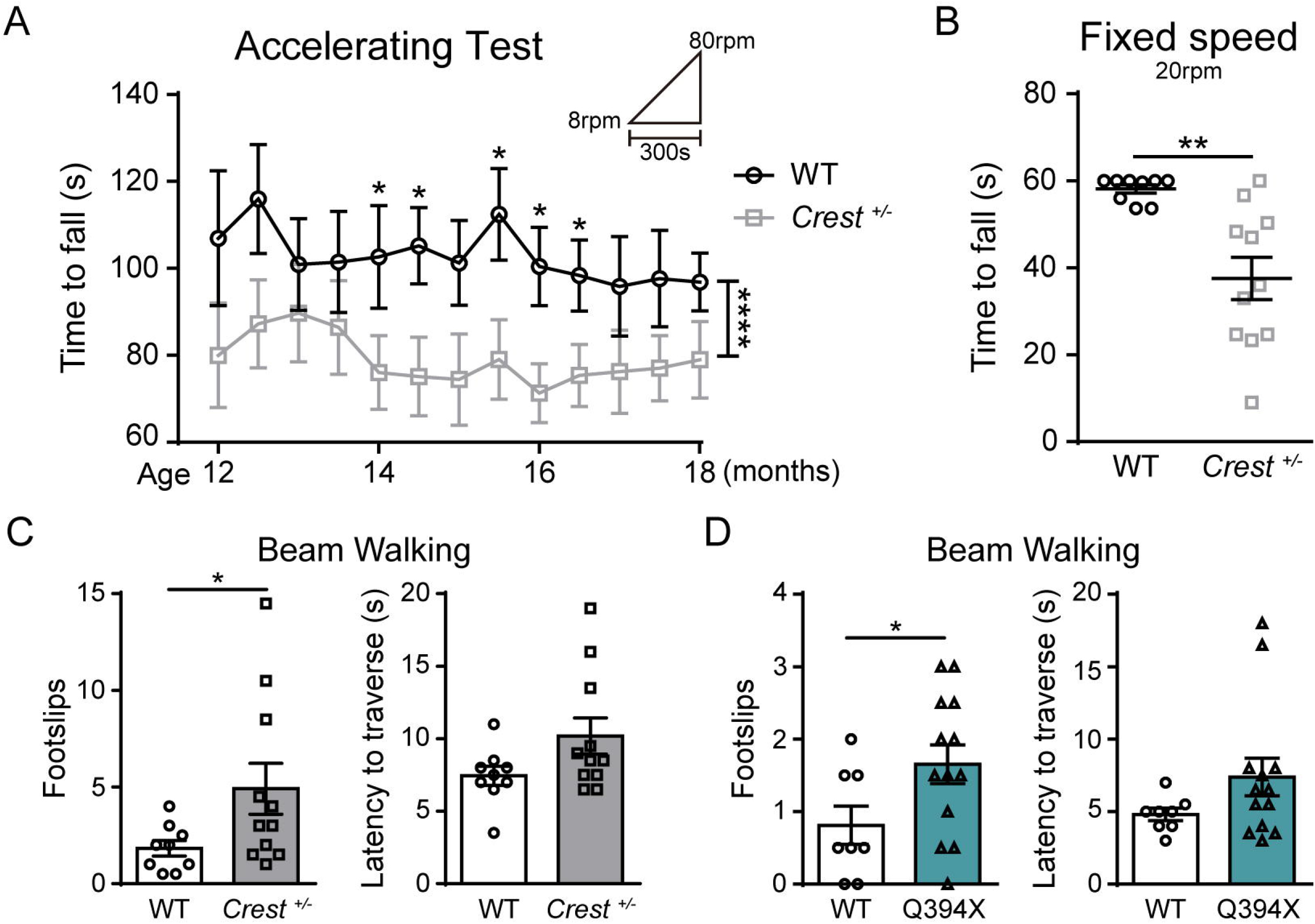
Both *CREST ^+/-^* and Q394X Mice Display Impaired Motor Coordination. (A) Behavioral performance of *Crest ^+/-^* mice (n=11) and WT littermates (n=9) in accelerating mode of rotarod tests from 8 rpm to 80 rpm within 300 seconds from the age of 12 months to 18 months. Tests were performed every two weeks. Error bars represent SEM. \**p* <0.05, Student’s *t* test; \*\*\*\**p* <0.0001, two-way ANOVA. (B) Performance of *Crest ^+/-^* mice (n=11) and WT littermates (n=9) in fixed speed mode of rotarod tests at 20 rpm up to 60 seconds at the age of 18 months. (C and D) The beam walking tests with square beam on *Crest ^+/-^* mice (n=11) and WT littermates (n=9) (C), and on Q394X mice (n=13) and WT littermates (n=8) (D) at 18 months of age. The performance was measured by times of hind paw slips (left panel) and the latency to traverse the beam (right panel). Error bars represent SEM. \**p* <0.05, \*\**p* < 0.01, Student’s *t* test.

We further examined the motor phenotypes of Q394X mice at 18 months of age. The general body weight and health of Q394X mice were also normal (Figure S4A). Although exhibiting no defects in open field and rotarod tests (Figures S4B and S4C), Q394X mice also showed more foot slips in the beam walking test compared to WT littermates (Figure 4D, Videos S3 and S4), indicating similar impairment of motor phenotypes in Q394X mice as in *Crest ^+/-^* mice. Taken together, we demonstrate that both haploinsufficiency and Q394X mutation of CREST can lead to elevated inflammatory responses in CNS and deficits in motor coordination *in vivo*.

### The CREST-BRG1 Complex Suppresses Expression of Cytokine Genes *Ccl2* and *Cxcl10* via Histone Deacetylation in Neurons

Prior to the investigation of molecular mechanisms underlying the increase of inflammatory responses in CREST deficient mice, we sought to determine the expression pattern of CREST in CNS. According to the data from Brain RNA-Seq database (Zhang et al., 2014) and previous report, *CREST* is highly expressed in neurons compared to other cell types in both human and mouse CNS (Figure S5A) (Aizawa et al., 2004). We confirmed this conclusion by performing immunohistochemistry experiments with various antibodies in cerebral motor cortex and lumbar spinal cord of adult WT mice. In agreement with previous data, we found that CREST mostly was not expressed in Iba1-positive microglia either in motor cortex or lumbar spinal cord (Figures S5B and S5C). However, CREST showed some expression in GFAP-positive astrocytes in white matter of lumbar spinal cord but not in motor cortex of WT mice (Figures S5B and S5C). Therefore, we suggest that neuronal loss of CREST may contribute to the microglial activation in CNS.

Given the fact that CREST is a critical transcriptional regulator (Qiu and Ghosh, 2008), we examined gene expression profiles by performing microarray analysis in primary cortical neurons infected with lentivirus harboring shRNA targeting CREST or control shRNA (Figures 5A, S6A and S6B). Using DAVID bioinformatics resources online (https://david.ncifcrf.gov) (Huang et al., 2009a, b), we performed gene ontology (GO) analysis of the microarray data and surprisingly found that the most significantly affected genes were predominantly associated with immune-associated processes (Figure 5B), suggesting the role of CREST in regulating the expression of genes related to inflammatory responses. Among the top 10 of the most remarkably upregulated genes, we selected two important chemokines, *Ccl2* and *Cxcl10*, which have been reported to play critical roles in microglial activation and inflammatory response, for further analysis (Clarner et al., 2015; Selenica et al., 2013). We first confirmed that mRNA levels of *Ccl2* and *Cxcl10* were significantly upregulated in CREST knockdown neurons compared to control neurons with real-time PCR (Figure 5C). Moreover, we observed a significant decrease in mRNA level of *Cxcl10* in CREST overexpressing neurons (Figures S6C and S6D). We then examined expression levels of endogenous *Ccl2* and *Cxcl10* in primary cortical neurons isolated from embryonic cortices of *Crest ^-/-^*, Q394X and WT mice, respectively. Importantly, we found that mRNA levels of both *Ccl2* and *Cxcl10* were markedly higher in *Crest ^-/-^* and Q394X neurons compared to WT littermates, respectively (Figures 5D and 5E). Thus, we demonstrate that neuronal loss of CREST leads to the altered expression of immune-related genes, such as the upregulation of chemokines *Ccl2* and *Cxcl10*, which may contribute to the microglial activation and elevated inflammatory responses.

**Figure 5.**
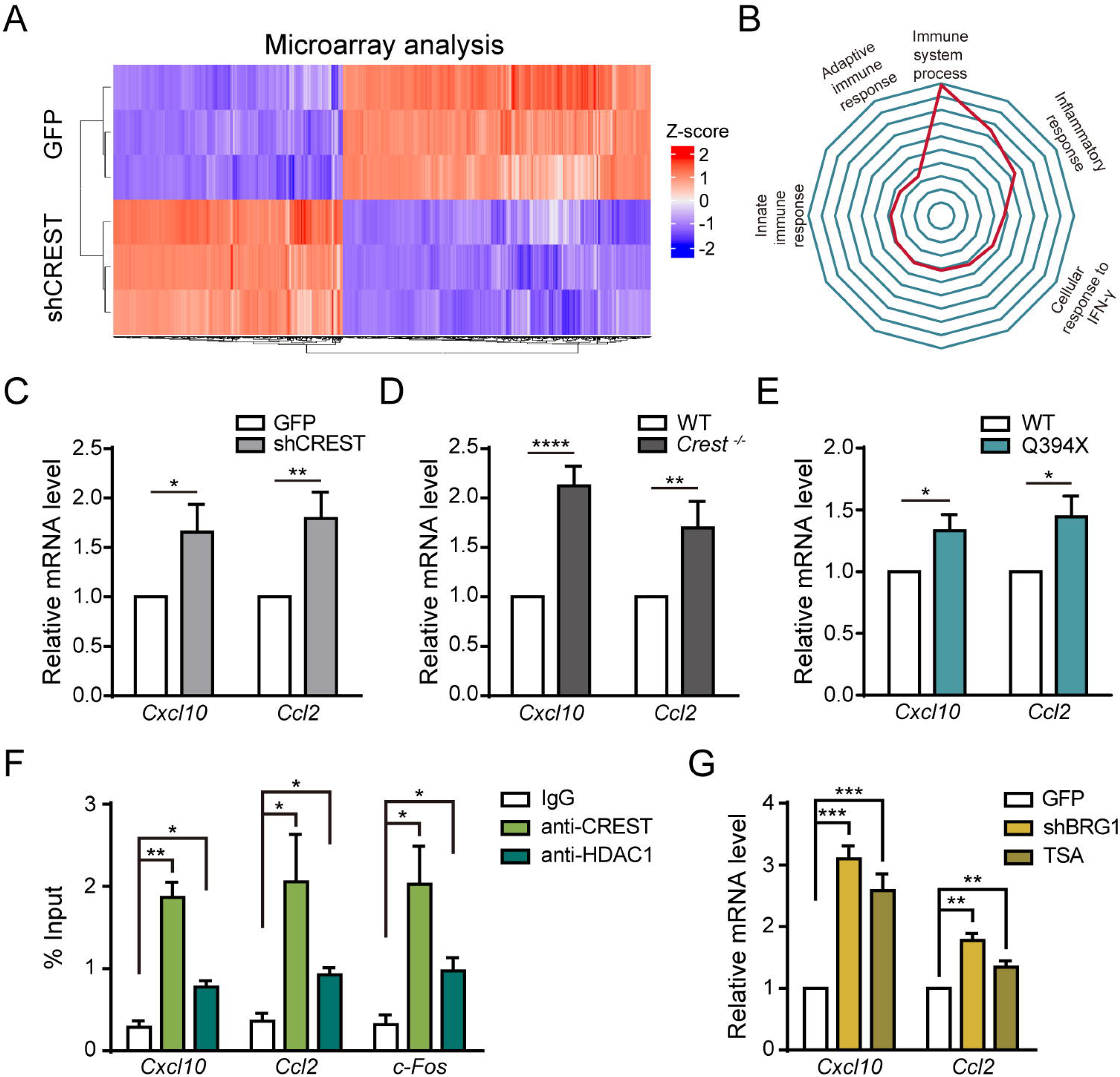
The CREST-BRG1 Complex Represses the Expression of Chemokine Genes via Recruiting Histone Deacetylation Complex in Neurons. (A) Heatmap showing the differentially expressed genes in primary cortical neurons infected with lentivirus expressing short hairpin RNA targeting CREST (shCREST) or expressing GFP (as control) by microarray analysis (fold change over 2.0). (B) Radar map showing the gene ontology analysis of gene lists in (A). (C) Quantitative RT-PCR of two chemokine genes, *Cxcl10* and *Ccl2*, for the confirmation of microarray data in primary cortical neurons transfected with shCREST or GFP as control by lentivirus. (D and E) Quantitative RT-PCR experiments validating the relative mRNA levels of *Cxcl10* and *Ccl2* in primary cortical neurons isolated from embryotic *Crest ^-/-^* (D) and Q394X (E) mice compared to their WT littermates, respectively. (F) Chromatin immunoprecipitation on promoter regions of *Cxcl10*, *Ccl2* and *c-Fos* genes with IgG, anti-CREST or anti-HDAC1 antibody in primary cortical neurons. Signals were normalized as percentage of input. (G) Quantitative RT-PCR of *Cxcl10* and *Ccl2* in primary cortical neurons treated with 0.5μM trichostatin A (TSA) for 6h, or infected by lentivirus expressing shRNA targeting BRG1 (shBRG1) or GFP (as control). Error bars represent SEM. \**p* <0.05, \*\**p* < 0.01, \*\*\**p* < 0.001, and \*\*\*\**p* <0.0001, Student’s *t* test.

To further reveal the molecular mechanism underlying the regulation of *Ccl2* and *Cxcl10* in neurons by CREST, we hypothesized that CREST inhibited the transcription of these two genes through the BRG1-CREST complex recruiting the histone deacetylase such as HDAC1 (Qiu and Ghosh, 2008). To test this hypothesis, we performed chromatin immunoprecipitation (ChIP) experiments in cultured neurons with anti-CREST and anti-HDAC1 antibodies. Indeed, we found that both CREST and HDAC1 interacted with the promoter regions of *Ccl2* and *Cxcl10* genes (Figure 5F). Moreover, mRNA levels of *Ccl2* and *Cxcl10* were increased in BRG1 knockdown cortical neurons by shRNA transfection (Figures S6E and S6F) compared to control neurons (Figure 5G). Consistently, we detected significantly upregulated mRNA levels of *Ccl2* and *Cxcl10* in neurons treated with an HDAC inhibitor trichostatin A (TSA) compared to controls (Figure 5G). Taken together, we suggest that the CREST-BRG1 complex plays a critical role in inhibiting the transcription of *Ccl2* and *Cxcl10* in neurons via HDAC-dependent histone deacetylation.

### Decreased Protein Level of FUS in the CNS of CREST KO Mice

Besides the regulation of immune-related genes by CREST in neurons, we also wonder whether CREST can influence the expression of other important ALS-linked genes, such as FUS which has been reported as a CREST-binding protein (Chesi et al., 2013) and TDP43. We performed immunohistochemistry with anti-FUS antibody in cerebral motor cortex and lumbar spinal cord sections of *Crest ^-/-^*, *Crest ^+/-^* and WT littermates at P14. Interestingly, we found that the level of FUS protein was stringently correlated with CREST protein level, and gradually decreased in both motor cortex and lumbar spinal cord of *Crest ^+/-^* and *Crest ^-/-^* mice compared to WT littermates (Figures 6A and 6B). Furthermore, we found that protein level of FUS rather than TDP43 was decreased in both frontal cortices and lumbar spinal cords collected from *Crest ^+/-^* mice compared to WT littermates at 18 months of age (Figures 6C and 6D). These data suggest that FUS appears to be *loss-of-function* in CREST mutant mice, further suggesting a converged pathway for ALS pathogenesis from different genetic causes.

**Figure 6.**
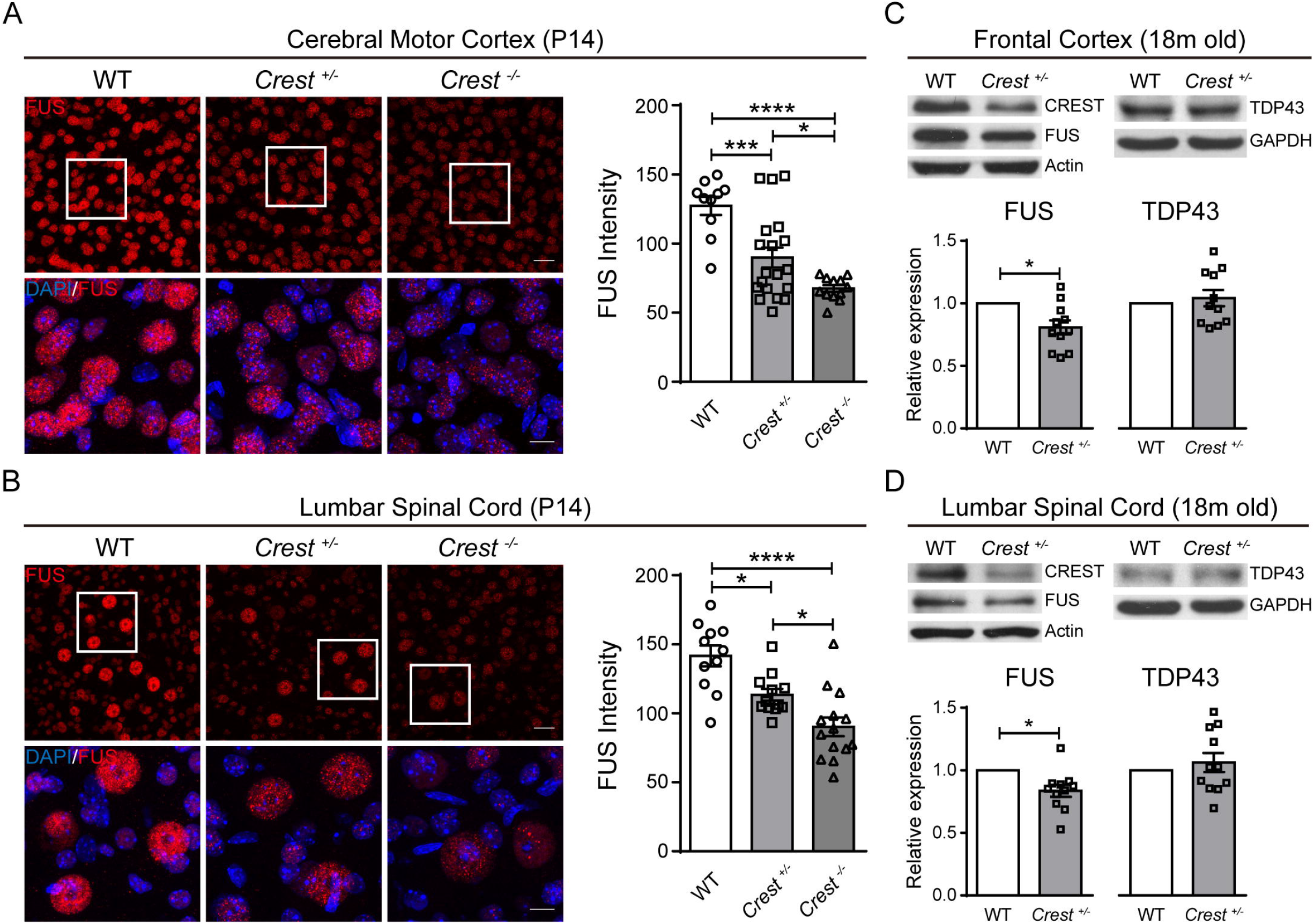
CREST and FUS Expression Represent Potentially Positive Correlation in CNS. (A and B) Representative confocal immunohistochemistry images (left panels) and intensity quantification (right graphs) of FUS (red) in the cerebral motor cortices (A) and the lumbar spinal cords (B) of *Crest ^-/-^* (n=3), *Crest ^+/-^* mice (n=3) and WT littermates (n=3) at P14. Top panels show lower magnification, scale bar = 25 μm. Bottom panels show higher magnification of the white square regions of upper panels, scale bar = 10 μm. Error bars represent SEM. \**p* <0.05, \*\*\**p* < 0.001, and \*\*\*\**p* <0.0001, one-way ANOVA. (C and D) Immunoblot (top panels) and intensity quantification (bottom graphs) of protein from the frontal cortices (C) and the lumbar spinal cords (D) of *Crest ^+/-^* mice (n=11) and WT littermates (n=5) at 18 months of age. Relative expression represents the normalized ratios of FUS/Actin (bottom left graphs) or of TDP43/GAPDH (bottom right graphs) band intensities determined by densitometry in Image J. Error bars represent SEM. *p<0.05, Student’s *t* test.

## DISCUSSION

Numerous publications have illustrated that chronic proinflammatory responses contribute to the pathogenic mechanisms underlying the progression of neurodegenerative diseases. In this study, we reveal that microglial appearance exhibits activation morphology including the enlargement of cell bodies and the decreased complexity of branches in both CREST KO and Q394X mutant mice. Since the differentiated subsets of activated microglia, including proinflammatory (M1) and antiinflammatory (M2) microglia (Kigerl et al., 2009), we detected the expression of some inflammatory genes in *Crest ^+/-^* mice and WT littermates at 18 months and found a specific upregulation of proinflammatory gene *Cox-2*, suggesting the overall effects of CREST haploinsufficiency on the chronic activation of neurotoxic M1 microglia, which may cause the axonal degeneration of motor neurons in the spinal cords. Although the increased expression of *Cox-2* in ALS patients and SOD1 mutant mice has been reported (Almer et al., 2001), one more interesting discovery is that the expression of *Cox-2* is specifically increased in ALS patients rather than other neurodegenerative disease cases (McGeer and McGeer, 2002). Moreover, inhibition of the enzymatic activity of COX-2 may delay the disease onset and ameliorate the survival of ALS mice (Drachman et al., 2002; Pompl et al., 2003), suggesting that chronic activation of M1 microglia and concomitant upregulation of *Cox-2* caused by loss of CREST contribute to ALS pathogenesis.

Consistent with axonal degeneration in CREST haploinsufficiency mice, we found that both *Crest ^+/-^* and Q394X mice exhibited deficits in motor coordination tasks, further confirming the connection between CREST and ALS pathogenesis. However, we did not detect very serious neurodegenerative phenotypes as in SOD1 mutant transgenic mice. Considering the involvement of both genetic and environmental factors in the pathogenesis of ALS, we may try some artificial manipulations, such as the administration of lipopolysaccharide (LPS), in CREST KO or mutant mice to induce more significant inflammatory responses and anticipate the appearance of more typical ALS symptoms (Nguyen et al., 2004).

We provide evidence to illustrate the alteration of immune-related genes, such as the upregulation of chemokines *Cxcl10* and *Ccl2*, in CREST deficient or ALS-related mutation Q388X-expressing neurons, which may play a critical role in microglial activation and increased proinflammatory responses *in vivo*. Moreover, we demonstrate that CREST may directly inhibit the transcription of *Cxcl10* and *Ccl2* genes through the CREST-BRG1-HDAC1 complex interacting with the promoter regions (Qiu and Ghosh, 2008). Consistent with the notion that upregulation of chemokines in CNS may contribute to the infiltration of periphery immune cells, increased expression of *Ccl2* in SOD1 mutant transgenic mice or ALS patients has been reported to induce the infiltration of dendritic cells and promote the acquisition of properties of antigen-presenting cells by microglia (Henkel et al., 2006; Henkel et al., 2004; Philips and Robberecht, 2011), suggesting the involvement of T cell infiltration in microglial activation in CREST mutant mice.

Another interesting pathological phenotype we observed in CREST KO mice is the decreased protein level of FUS in CNS. Previous publications have shown that knockdown of FUS can reduce the cell viability in vitro, and cause the anatomical loss of neuromuscular junctions and concomitant impairment of locomotion ability in Drosophila and zebrafish models (Armstrong and Drapeau, 2013; Sasayama et al., 2012; Ward et al., 2014), suggesting that besides the neurotoxic proinflammation induced by microglial activation, the loss of FUS may also contribute to the degeneration of motor neurons in CREST KO mice. In conclusion, we demonstrate that *CREST* is a pathogenic gene for ALS, mutations of which lead to upregulation of chemokine genes in neurons, subsequently activate microglia, increase proinflammatory responses *in vivo*, and ultimately impair motor function in aged CREST KO and Q394X mutant mice. These data provide a transcriptional pathway for neuroinflammation activation in ALS, suggesting that approaches of modulating histone deacetylase complex may be the candidates for therapeutic intervenes.

## STAR METHODS

Detailed methods are provided in the online version for this paper and include the following:

## KEY RESOURCES TABLE

**CONTACT FOR REAGENT AND RESOURCE SHARING**

**EXPERIMENTAL MODEL AND SUBJECT DETAILS**

## METHOD DETAILS

Plasmid construction

Animals and ethics statement

Primary cortical neuron culture

Western blotting

Immunohistochemistry

Morphological analysis

RNA isolation and quantitative RT-PCR

Chromatin immunoprecipitation

Behavior tests

## QUANTIFICATION AND STATISTICAL ANALYSIS

### SUPPLEMENTAL INFORMATION

Supplemental Information includes six figures and four videos.

## ACKNOWLEDGMENTS

We thank ION transgenic core facility for making CREST knockout and Q394X mutant mice in this work. This work was supported by NSFC Grants (#31625013, #91732302) and Shanghai Brain-Intelligence Project from STCSM (16JC1420501). The research is supported by the Open Large Infrastructure Research of Chinese Academy of Sciences. The authors declare that they do not have any conflicts of interest for this work.

## AUTHOR CONTRIBUTIONS

Z.Q. conceived and supervised the project. C.C. performed most parts of experiments and data analysis. K.Y. and X.W. performed parts of biochemistry experiments. Y.Z. managed mice for experiments. S.S. isolated primary neurons. A.Ghosh contributed to conditional CREST mice construction. A.Gitler contributed to the genetic data containing CREST Q388X mutations. Z.Q. wrote the manuscript.

---

**Figure SI.**
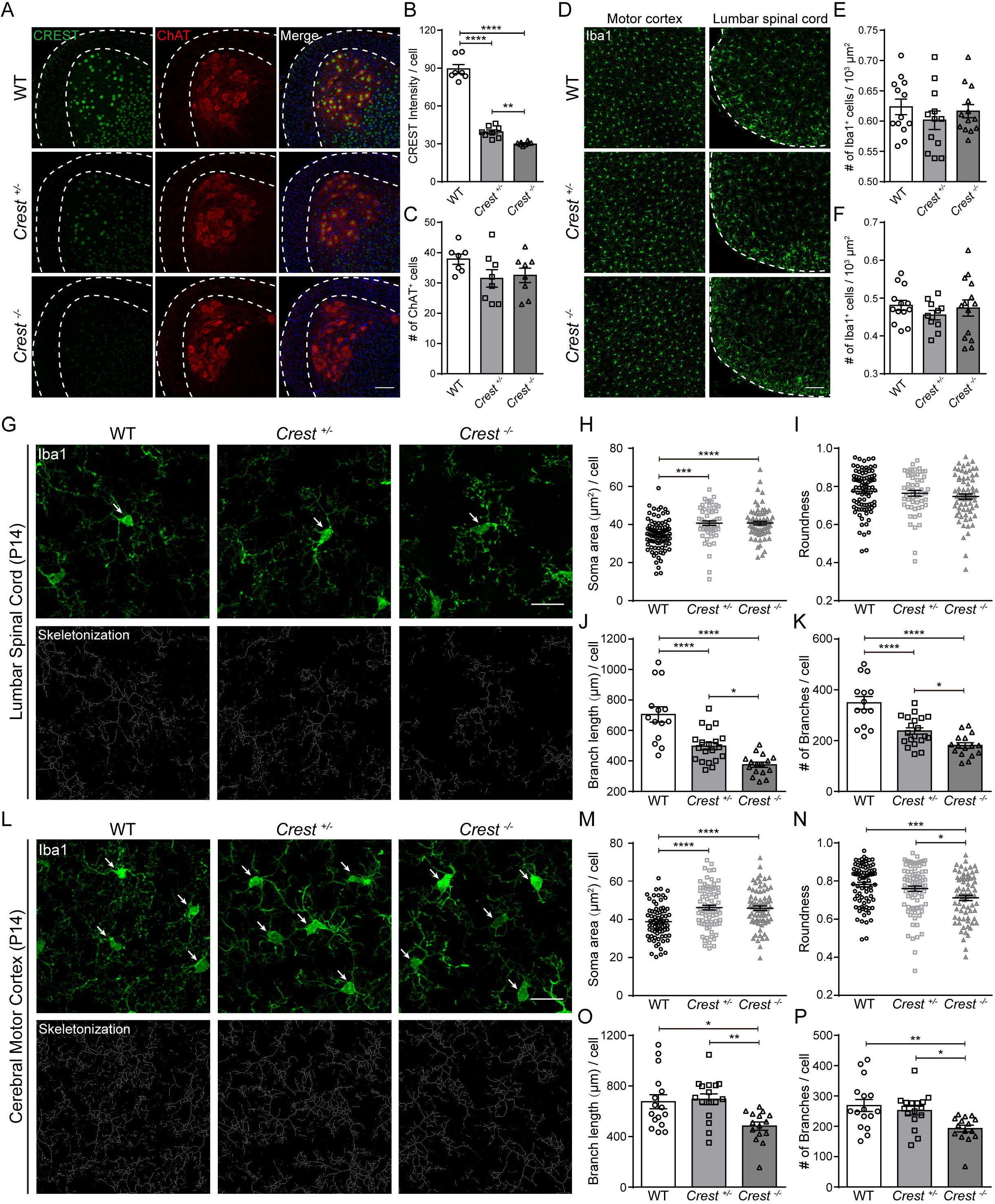
CREST KO Mice Show the Activation of Microglia in CNS at P14, related to Figure 2. (A) Representative confocal immunohistochemistry images of CREST-positive cells (green) and ChAT-positive motor neurons (red) in the lumbar spinal cords of P14 *Crest^-/-^* (n=3), *Crest^+/-^* mice (n=3) and WT littermates (n=3). Dashed lines divide the white and grey matter of lumbar spinal cord. Scale bar, 100μm. (B and C) Quantification of CREST intensities (B) and the number of ChAT-positive motor neurons (C) in the lumbar spinal cords of P14 *Crest^-/-^* (n=3), *Crest^+/-^* mice (n=3) and WT littermates (n=3). (D) Representative confocal immunohistochemistry images of Ibal-positive microglia (green) in cerebral motor cortices (left panels) and in lumbar spinal cords (right panels) of P14 *Crest*(n=3), *Crest^+/-^* mice (n=3) and WT littermates (n=3). Dashed lines show the lumbar spinal cords. Scale bar, 100μm. (E and F) Quantification of Ibal-positive microglia (green) in cerebral motor cortices (E) and in lumbar spinal cords (F) of P14 *Crest^-/-^* (n=3), *Crest^+/-^* mice (n=3) and WT littermates (n=3). (G) Representative immunohistochemistry images of Ibal-positive (green; arrows) microglia (top panels) and their skeletonized appearance (bottom panels) in the lumbar spinal cords of P14 Cres(n=3), *Crest^+/-^* mice (n=3) and WT littermates (n=3). Scale bar, 20μm. (H-K) Quantification of microglial morphological parameters including soma area (H) and roundness (I) of the projection of Ibal-positive cell bodies, and branch length (J) and branch number (K) per cell in the lumbar spinal cords of P14 *Crest^-/-^* (n=3), *Crest^+/-^* mice (n=3) and WT littermates (n=3). (L) Representative imiuunohistochemistry images of Ibal-positive (green; arrows) microglia (top panels) and their skeletonized appearance (bottom panels) in the cerebral motor cortices of P14 *Crest^-/-^* (n=3), *Crest^+/-^* mice (n=3) and WT littermates (n=3). Scale bar, 20μm. (M-P) Quantification of microglial morphological parameters including soma area (M) and roundness (N) of the projection of Ibal-positive cell bodies, and branch length (O) and branch number (P) per cell in the cerebral motor cortices of P14 *Crest^-/-^* (n=3), *Crest^+/-^* mice (n=3) and WT littermates (n=3). Error bars represent SEM. *p<0.05, **p< 0.01, ***p< 0.001, and ****p<0.0001, one-way ANOVA.

**Figure S2.**
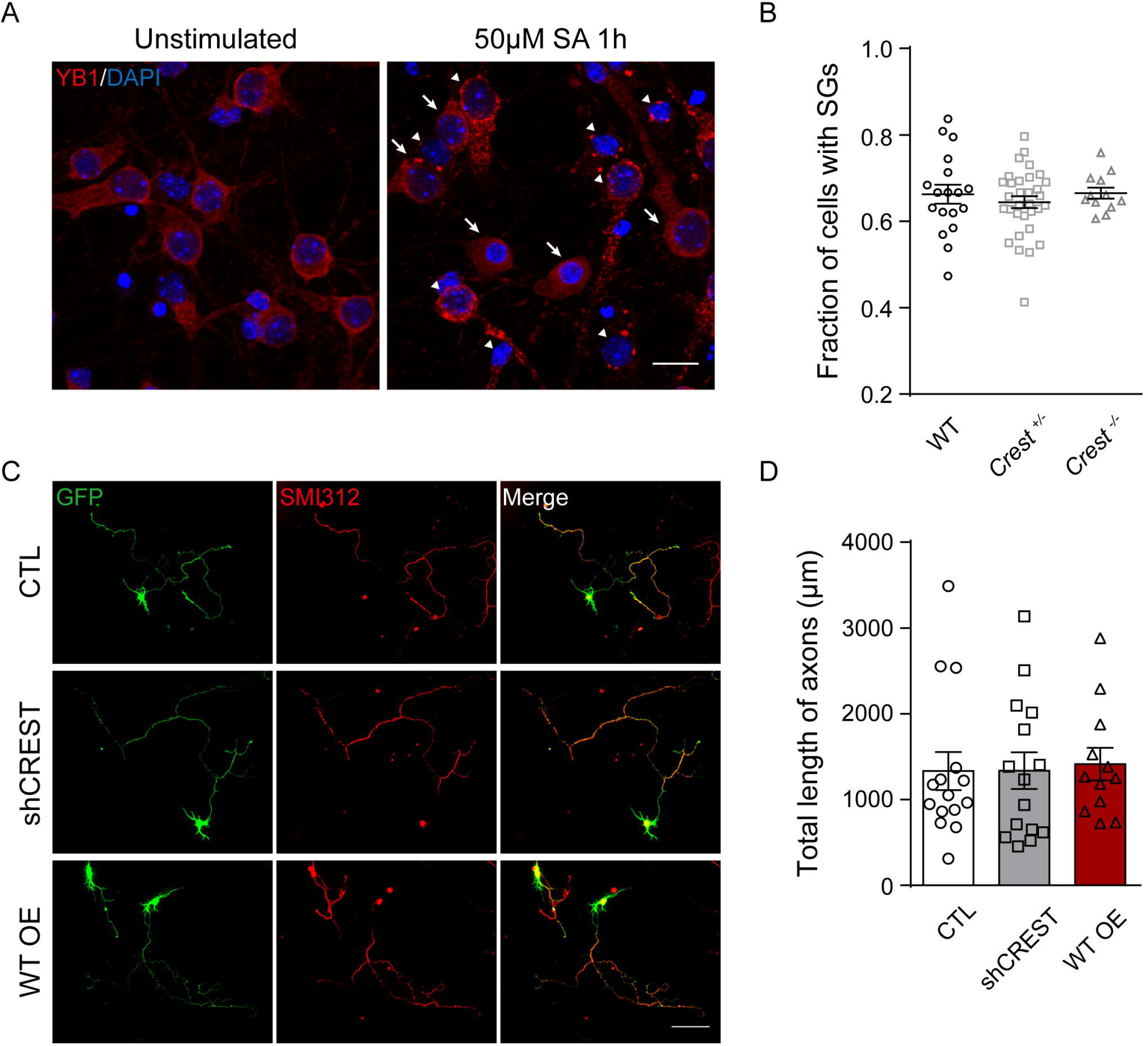
Loss of CREST Function Does not Affect the Formation of Stress Granules and the Axon Length in Cultured Neurons /7z *vitro*, related to Figure 2. (A) Representative Immunofluorescence staining of stress granule marker YB1 (red) in the treatment of nothing as control (left) or 50μΜ sodium arsenite (SA) for lh (right). Arrows indicate the cells without stress granules (SGs) in the treatment of SA. Arrowheads indicate the cells with SGs. Scale bar, 15μm. (B) Quantification of the fraction of cells with SGs in cultured neurons isolated from embryotic *Crest^-/-^, Crest^+/-^* mice and WT littermates. (C and D) Representative Immunofluorescence images (C) and the total length quantification of SMI312-positive (red) axons in cultured neurons expressing GFP (C, top), shCREST labeled by GFP (C, middle) and WT human CREST labeled by GFP (green) (C, bottom). Scale bar, 100μm. Error bars represent SEM. One-way ANOVA.

**Figure S3.**
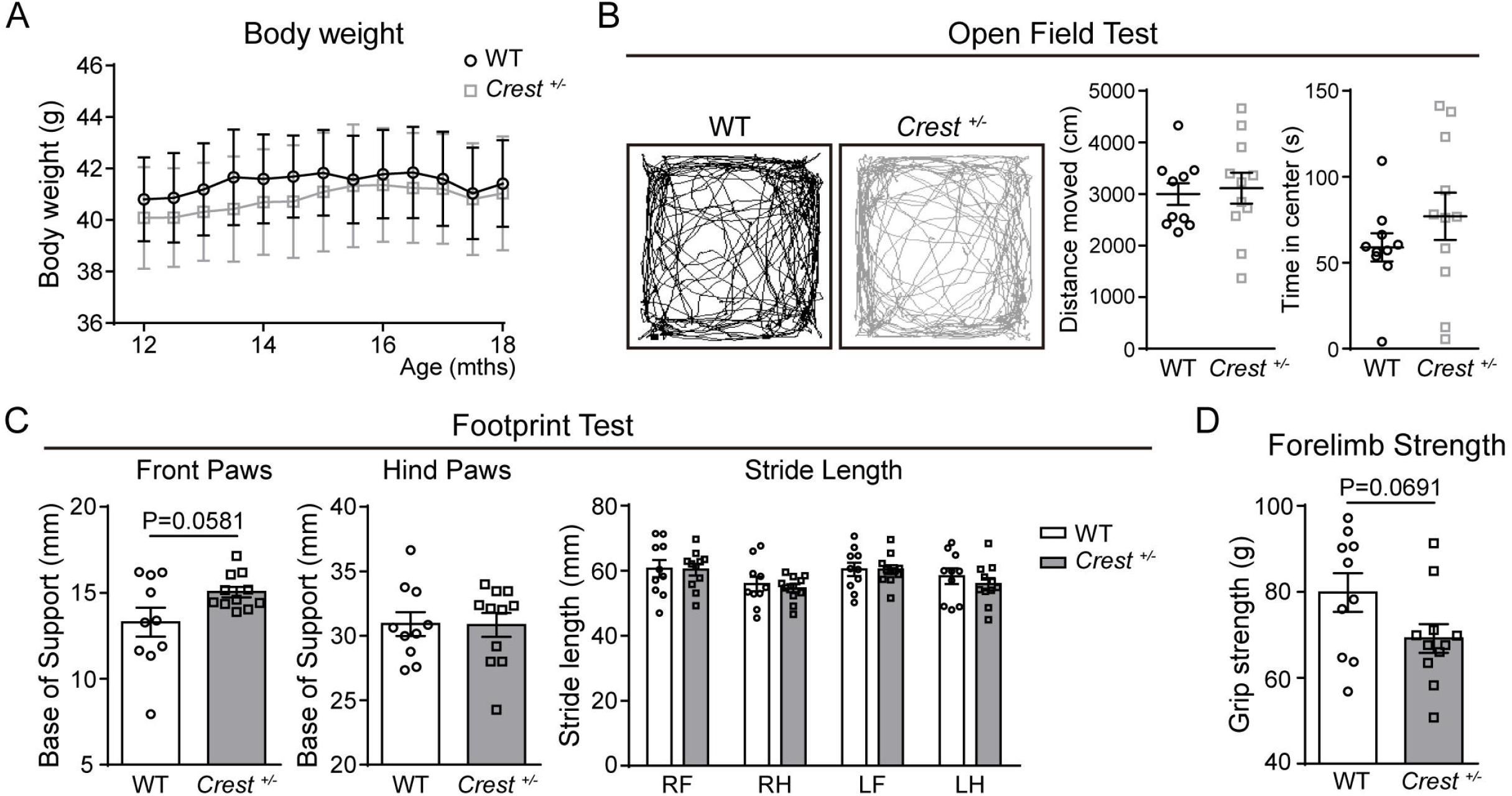
Behavioral Tests of Motor Phenotypes on *Crest^+/-^* Mice, related to Figure 4. (A) Body weight of *Crest^+/-^* mice (n=11) and WT littermates (n=9) from the age of 12 months to 18 months. The measurement was performed every 2 weeks. (B) Representative traces (two left panels), the moving distance (middle graph) and the center staying time (right graph) of *Crest^+/-^* mice (n=11) and WT littermates (n=9) at 18 months in open field tests. The duration of each trial was 10 min. (C) Footprint tests showing the width of support bases of both front paws (left graph) and hind paws (middle graph), and the length of strides of four paws (right graph) of *Crest^+/-^* mice (n=11) and WT littermates (n=9) at 18 months. (D) Grip strength tests showing the forelimb strength of *Crest^+/-^* mice (n=11) and WT littermates (n=9) at 18 months. Error bars represent SEM. Student’s *t* test.

**Figure S4.**
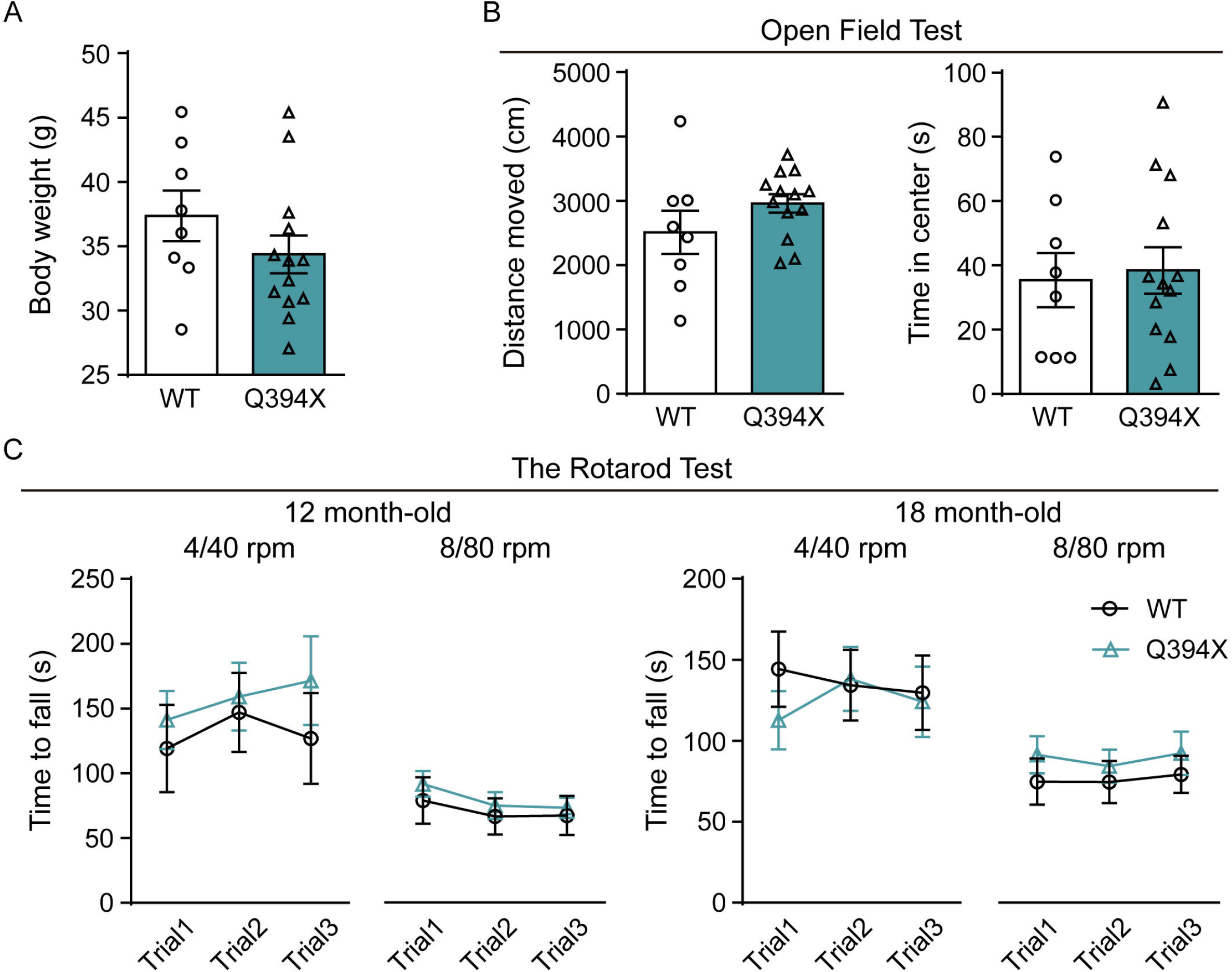
Behavioral Tests of Motor Phenotypes on Q394X Mice, related to Figure 4. (A) Body weight of Q394X mice (n=13) and WT littermates (n=8) at 18 months. (B) Open field tests showing the moving distance (left graph) and the center staying time (right graph of Q394X mice (n=13) and WT littermates (n=8) at 18 months. The duration of each trial was 10 min. (C) Rotarod tests performed with two accelerating modes (4 to 40 rpm and 8 to 80 rpm in 300 seconds) for 3 trials on Q394X mice (n=13) and WT littermates (n=8) at the age of 12 months (two left graphs) and 18 months (two right graphs). Error bars represent SEM. Student’s / test.

**Figure S5.**
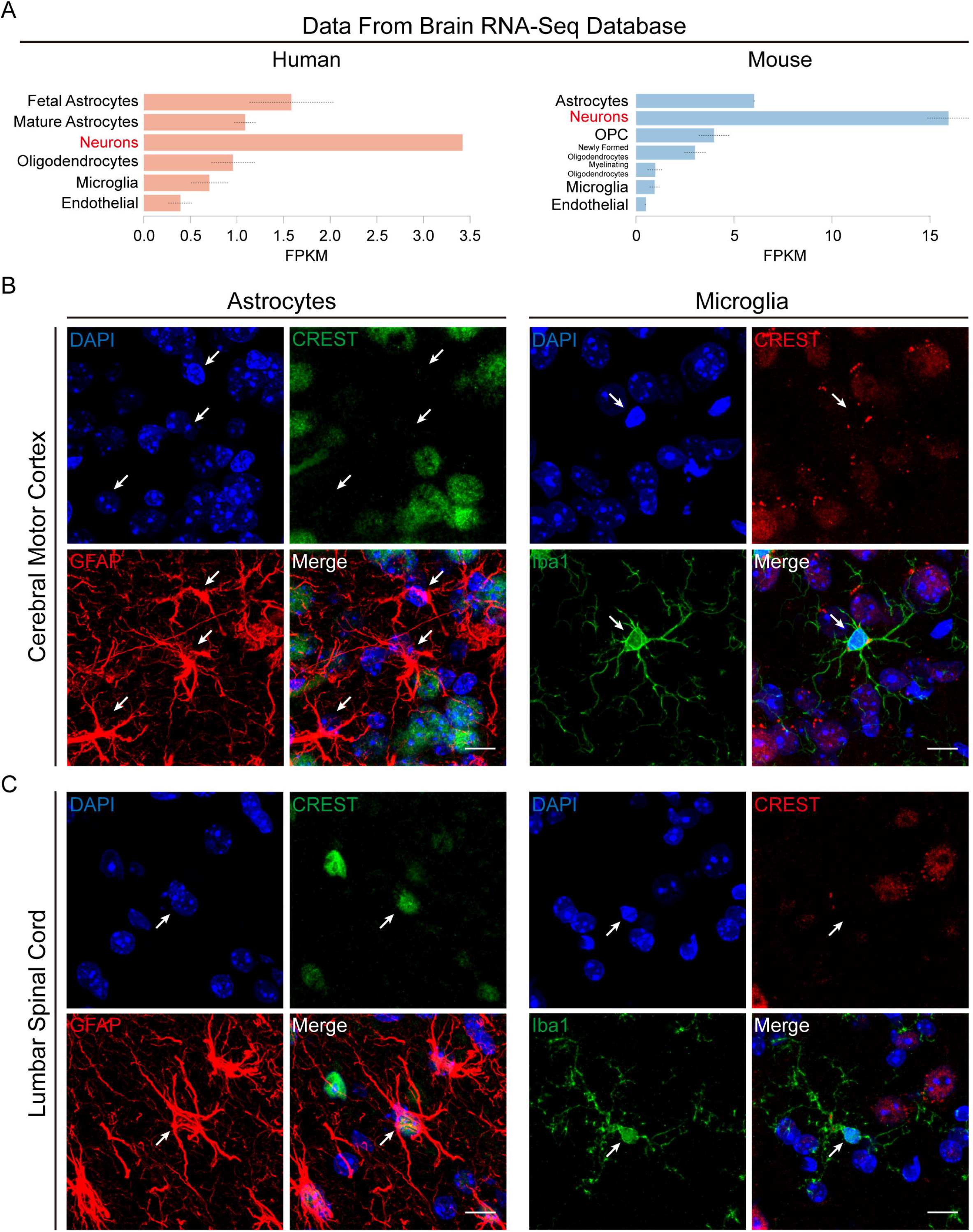
Expression Pattern of CREST in the CNS, related to Figure 5. (A) Relative expression of CREST in cell types of human (left) and mouse (right) brain revealed by Brain RNA-Seq database. (B and C) Representative confocal immunohistochemistry images of co-localization of astrocyte marker GFAP (red) and CREST (green) (four left panels), and of microglial marker Ibal (green) and CREST (red) (four right panels) in both cerebral motor cortex (B) and lumbar spinal cord (C). Arrows indicate GFAP-positive astrocytes or Ibal-positive microglia. Scale bar, 10μm.

**Figure S6.**
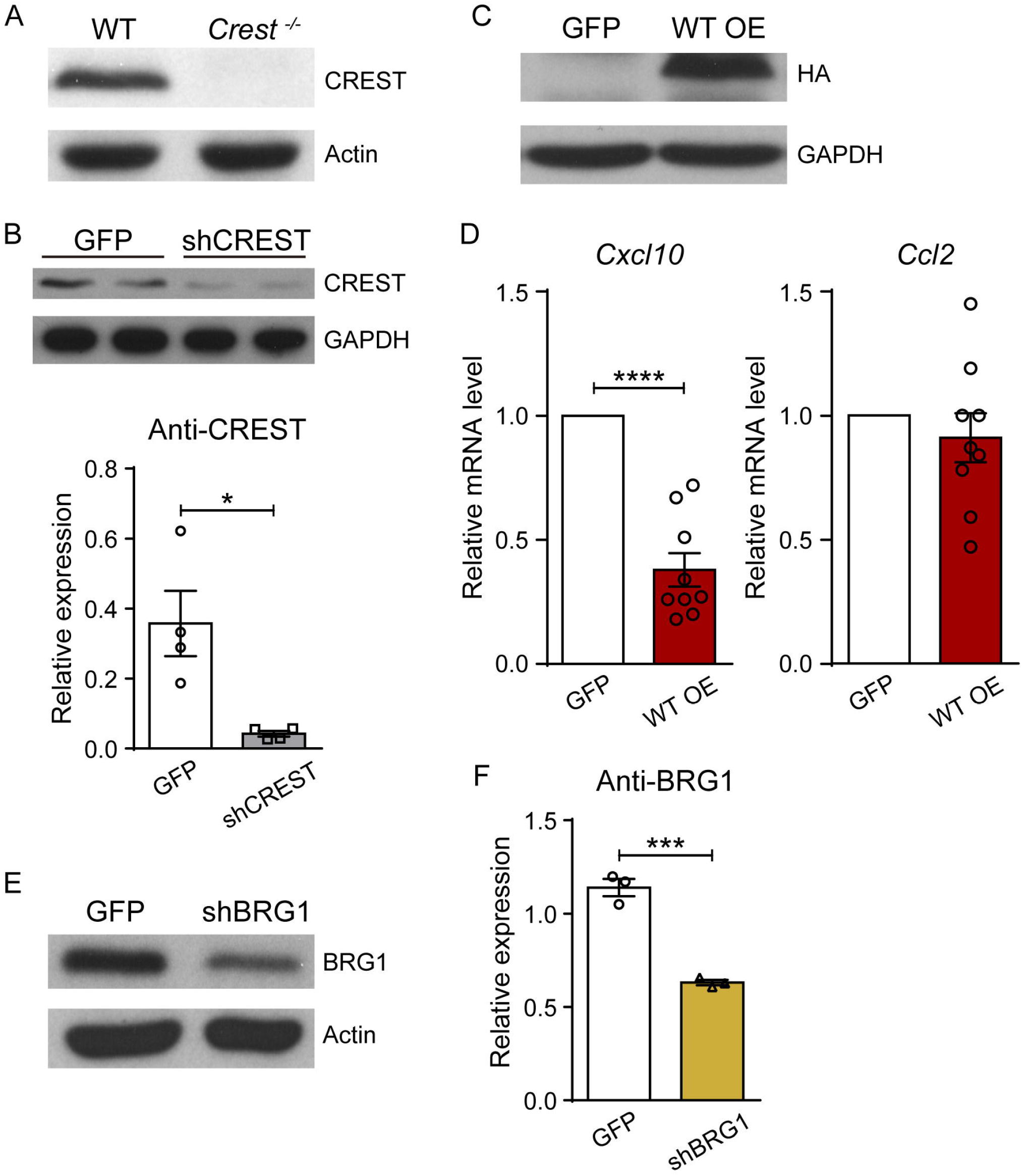
Validation of Knockdown Efficiency of shRNAs and the Inhibitory Effects of CREST on the Transcription of *Cxcl10* in Cultured Neurons, related to Figure 5. (A) Immunoblot of protein from primary cortical neurons isolated from embryotic *Crest^-/-^* and WT littermates, using anti-CREST (top) and anti-Actin (bottom) antibodies. (B) Immunoblot and intensity quantification of protein from primary cortical neurons infected with lentivirus carrying CREST shRNA (shCREST) and GFP (as control). Relative expression represents the intensity ratios of CREST/GAPDH. (C) Immunoblot of protein from primary cortical neurons infected with lentivirus expressing GFP or HA-tagged WT CREST, using anti-HA (top) and anti-GAPDH (bottom) antibodies. (D) Quantitative RT-PCR of *Cxcl10* (left) and *Ccl2* (right) in primary cortical neurons infected with lentivirus expressing GFP or HA-tagged WT CREST. (E and F) Immunoblot (E) and quantification (F) of protein from primary cortical neurons infected with lentivirus carrying BRG1 shRNA (shBRGl) and GFP (as control). Relative expression represents the intensity ratios of BRGl/Actin normalized to one control sample. Error bars represent SEM. *p* <0.05, \*\*\*\**p* < 0.001, and <0.0001, Student’s /test.

## REFERENCES

Aizawa, H., Hu, S.C., Bobb, K., Balakrishnan, K., Ince, G., Gurevich, I., Cowan, M., and Ghosh, A. (2004). Dendrite development regulated by CREST, a calcium-regulated transcriptional activator. Science 303, 197–202.

Alexianu, M.E., Kozovska, M., and Appel, S.H. (2001). Immune reactivity in a mouse model of familial ALS correlates with disease progression. Neurology 57, 1282–1289.

Almer, G., Guegan, C., Teismann, P., Naini, A., Rosoklija, G., Hays, A.P., Chen, C., and Przedborski, S. (2001). Increased expression of the pro-inflammatory enzyme cyclooxygenase-2 in amyotrophic lateral sclerosis. Annals of neurology 49, 176–185.

Armstrong, G.A.B., and Drapeau, P. (2013). Loss and gain of FUS function impair neuromuscular synaptic transmission in a genetic model of ALS. Human molecular genetics 22, 4282–4292.

Beers, D.R., Zhao, W.H., Liao, B., Kano, O., Wang, J.H., Huang, A.L., Appel, S.H., and Henkel, J.S. (2011). Neuroinflammation modulates distinct regional and temporal clinical responses in ALS mice. Brain Behav Immun 25, 1025–1035.

Boillee, S., Yamanaka, K., Lobsiger, C.S., Copeland, N.G., Jenkins, N.A., Kassiotis, G., Kollias, G., and Cleveland, D.W. (2006). Onset and progression in inherited ALS determined by motor neurons and microglia. Science 312, 1389–1392.

Brooks, S.P., and Dunnett, S.B. (2009). Tests to assess motor phenotype in mice: a user’s guide. Nat Rev Neurosci 10, 519–529.

Carter, R.J., Morton, J., and Dunnett, S.B. (2001). Motor coordination and balance in rodents. Current protocols in neuroscience Chapter 8, Unit 8 12.

Chesi, A., Staahl, B.T., Jovicic, A., Couthouis, J., Fasolino, M., Raphael, A.R., Yamazaki, T., Elias, L., Polak, M., Kelly, C., et al. (2013). Exome sequencing to identify de novo mutations in sporadic ALS trios. Nature neuroscience 16, 851–855.

Cirulli, E.T., Lasseigne, B.N., Petrovski, S., Sapp, P.C., Dion, P.A., Leblond, C.S., Couthouis, J., Lu, Y.F., Wang, Q.L., Krueger, B.J., et al. (2015). Exome sequencing in amyotrophic lateral sclerosis identifies risk genes and pathways. Science 347, 1436–1441.

Clarner, T., Janssen, K., Nellessen, L., Stangel, M., Skripuletz, T., Krauspe, B., Hess, F.M., Denecke, B., Beutner, C., Linnartz-Gerlach, B., et al. (2015). CXCL10 triggers early microglial activation in the cuprizone model. Journal of immunology 194, 3400–3413.

Clement, A.M., Nguyen, M.D., Roberts, E.A., Garcia, M.L., Boillee, S., Rule, M., McMahon, A.P., Doucette, W., Siwek, D., Ferrante, R.J., et al. (2003). Wild-type nonneuronal cells extend survival of SOD1 mutant motor neurons in ALS mice. Science 302, 113–117.

DeJesus-Hernandez, M., Mackenzie, I.R., Boeve, B.F., Boxer, A.L., Baker, M., Rutherford, N.J., Nicholson, A.M., Finch, N.A., Flynn, H., Adamson, J., et al. (2011). Expanded GGGGCC hexanucleotide repeat in noncoding region of C9ORF72 causes chromosome 9p-linked FTD and ALS. Neuron 72, 245–256.

Drachman, D.B., Frank, K., Dykes-Hoberg, M., Teismann, P., Almer, G., Przedborski, S., and Rothstein, J.D. (2002). Cyclooxygenase 2 inhibition protects motor neurons and prolongs survival in a transgenic mouse model of ALS. Annals of neurology 52, 771–778.

Engelhardt, J.I., and Appel, S.H. (1990). Igg Reactivity in the Spinal-Cord and Motor Cortex in Amyotrophic-Lateral-Sclerosis. Arch Neurol-Chicago 47, 1210–1216.

Engelhardt, J.I., Tajti, J., and Appel, S.H. (1993). Lymphocytic Infiltrates in the Spinal-Cord in Amyotrophic-Lateral-Sclerosis. Arch Neurol-Chicago 50, 30–36.

Frakes, A.E., Ferraiuolo, L., Haidet-Phillips, A.M., Schmelzer, L., Braun, L., Miranda, C.J., Ladner, K.J., Bevan, A.K., Foust, K.D., Godbout, J.P., et al. (2014). Microglia induce motor neuron death via the classical NF-kappaB pathway in amyotrophic lateral sclerosis. Neuron 81, 1009–1023.

Glass, C.K., Saijo, K., Winner, B., Marchetto, M.C., and Gage, F.H. (2010). Mechanisms Underlying Inflammation in Neurodegeneration. Cell 140, 918–934.

Hall, E.D., Oostveen, J.A., and Gurney, M.E. (1998). Relationship of microglial and astrocytic activation to disease onset and progression in a transgenic model of familial ALS. Glia 23, 249–256.

Henkel, J.S., Beers, D.R., Siklos, L., and Appel, S.H. (2006). The chemokine MCP-1 and the dendritic and myeloid cells it attracts are increased in the mSOD1 mouse model of ALS. Molecular and cellular neurosciences 31, 427–437.

Henkel, J.S., Engelhardt, J.I., Siklos, L., Simpson, E.P., Kim, S.H., Pan, T., Goodman, J.C., Siddique, T., Beers, D.R., and Appel, S.H. (2004). Presence of dendritic cells, MCP-1, and activated microglia/macrophages in amyotrophic lateral sclerosis spinal cord tissue. Annals of neurology 55, 221–235.

Huang, D.W., Sherman, B.T., and Lempicki, R.A. (2009a). Bioinformatics enrichment tools: paths toward the comprehensive functional analysis of large gene lists. Nucleic Acids Res 37, 1–13.

Huang, D.W., Sherman, B.T., and Lempicki, R.A. (2009b). Systematic and integrative analysis of large gene lists using DAVID bioinformatics resources. Nat Protoc 4, 44–57.

Kawamata, T., Akiyama, H., Yamada, T., and McGeer, P.L. (1992). Immunologic reactions in amyotrophic lateral sclerosis brain and spinal cord tissue. The American journal of pathology 140, 691–707.

Kigerl, K.A., Gensel, J.C., Ankeny, D.P., Alexander, J.K., Donnelly, D.J., and Popovich, P.G. (2009). Identification of Two Distinct Macrophage Subsets with Divergent Effects Causing either Neurotoxicity or Regeneration in the Injured Mouse Spinal Cord. J Neurosci 29, 13435–13444.

Kukharsky, M.S., Quintiero, A., Matsumoto, T., Matsukawa, K., An, H., Hashimoto, T., Iwatsubo, T., Buchman, V.L., and Shelkovnikova, T.A. (2015). Calcium-responsive transactivator (CREST) protein shares a set of structural and functional traits with other proteins associated with amyotrophic lateral sclerosis. Molecular neurodegeneration 10, 20.

Kwiatkowski, T.J., Jr., Bosco, D.A., Leclerc, A.L., Tamrazian, E., Vanderburg, C.R., Russ, C., Davis, A., Gilchrist, J., Kasarskis, E.J., Munsat, T., et al. (2009). Mutations in the FUS/TLS gene on chromosome 16 cause familial amyotrophic lateral sclerosis. Science 323, 1205–1208.

Mackenzie, I.R.A., Bigio, E.H., Ince, P.G., Geser, F., Neumann, M., Cairns, N.J., Kwong, L.K., Forman, M.S., Ravits, J., Stewart, H., et al. (2007). Pathological TDP-43 distinguishes sporadic amyotrophic lateral sclerosis from amyotrophic lateral sclerosis with SOD1 mutations. Annals of neurology 61, 427–434.

Mantovani, S., Garbelli, S., Pasini, A., Alimonti, D., Perotti, C., Melazzini, M., Bendotti, C., and Mora, G. (2009). Immune system alterations in sporadic amyotrophic lateral sclerosis patients suggest an ongoing neuroinflammatory process. J Neuroimmunol 210, 73–79.

Mass, E., Jacome-Galarza, C.E., Blank, T., Lazarov, T., Durham, B.H., Ozkaya, N., Pastore, A., Schwabenland, M., Chung, Y.R., Rosenblum, M.K., et al. (2017). A somatic mutation in erythro-myeloid progenitors causes neurodegenerative disease. Nature 549, 389–+.

McGeer, P.L., and McGeer, E.G. (2002). Inflammatory processes in amyotrophic lateral sclerosis. Muscle & nerve 26, 459–470.

Meissner, F., Molawi, K., and Zychlinsky, A. (2010). Mutant superoxide dismutase 1-induced IL-1beta accelerates ALS pathogenesis. Proc Natl Acad Sci U S A 107, 13046–13050.

Morrison, H.W., and Filosa, J.A. (2013). A quantitative spatiotemporal analysis of microglia morphology during ischemic stroke and reperfusion. Journal of neuroinflammation 10.

Neumann, M., Sampathu, D.M., Kwong, L.K., Truax, A.C., Micsenyi, M.C., Chou, T.T., Bruce, J., Schuck, T., Grossman, M., Clark, C.M., et al. (2006). Ubiquitinated TDP-43 in frontotemporal lobar degeneration and amyotrophic lateral sclerosis. Science 314, 130–133.

Neymotin, A., Petri, S., Calingasan, N.Y., Wille, E., Schafer, P., Stewart, C., Hensley, K., Beal, M.F., and Kiaei, M. (2009). Lenalidomide (Revlimid (R)) administration at symptom onset is neuroprotective in a mouse model of amyotrophic lateral sclerosis. Exp Neurol 220, 191–197.

Nguyen, M.D., D’Aigle, T., Gowing, G., Julien, J.P., and Rivest, S. (2004). Exacerbation of motor neuron disease by chronic stimulation of innate immunity in a mouse model of amyotrophic lateral sclerosis. J Neurosci 24, 1340–1349.

Philips, T., and Robberecht, W. (2011). Neuroinflammation in amyotrophic lateral sclerosis: role of glial activation in motor neuron disease. The Lancet Neurology 10, 253–263.

Pompl, P.N., Ho, L., Bianchi, M., McManus, T., Qin, W., and Pasinetti, G.M. (2003). A therapeutic role for cyclooxygenase-2 inhibitors in a transgenic mouse model of amyotrophic lateral sclerosis. FASEB journal : official publication of the Federation of American Societies for Experimental Biology 17, 725–727.

Qiu, Z., and Ghosh, A. (2008). A Calcium-Dependent Switch in a CREST-BRG1 Complex Regulates Activity-Dependent Gene Expression. Neuron 60, 775–787.

Renton, A.E., Majounie, E., Waite, A., Simon-Sanchez, J., Rollinson, S., Gibbs, J.R., Schymick, J.C., Laaksovirta, H., van Swieten, J.C., Myllykangas, L., et al. (2011). A Hexanucleotide Repeat Expansion in C9ORF72 Is the Cause of Chromosome 9p21-Linked ALS-FTD. Neuron 72, 257–268.

Rosen, D.R., Siddique, T., Patterson, D., Figlewicz, D.A., Sapp, P., Hentati, A., Donaldson, D., Goto, J., O’Regan, J.P., Deng, H.X., and, et al. (1993). Mutations in Cu/Zn superoxide dismutase gene are associated with familial amyotrophic lateral sclerosis. Nature 362, 59–62.

Sasayama, H., Shimamura, M., Tokuda, T., Azuma, Y., Yoshida, T., Mizuno, T., Nakagawa, M., Fujikake, N., Nagai, Y., and Yamaguchi, M. (2012). Knockdown of the Drosophila Fused in Sarcoma (FUS) Homologue Causes Deficient Locomotive Behavior and Shortening of Motoneuron Terminal Branches. Plos One 7.

Selenica, M.L., Alvarez, J.A., Nash, K.R., Lee, D.C., Cao, C., Lin, X., Reid, P., Mouton, P.R., Morgan, D., and Gordon, M.N. (2013). Diverse activation of microglia by chemokine (C-C motif) ligand 2 overexpression in brain. Journal of neuroinflammation 10, 86.

Sreedharan, J., Blair, I.P., Tripathi, V.B., Hu, X., Vance, C., Rogelj, B., Ackerley, S., Durnall, J.C., Williams, K.L., Buratti, E., et al. (2008). TDP-43 mutations in familial and sporadic amyotrophic lateral sclerosis. Science 319, 1668–1672.

Taylor, J.P., Brown, R.H., Jr., and Cleveland, D.W. (2016). Decoding ALS: from genes to mechanism. Nature 539, 197–206.

Teyssou, E., Vandenberghe, N., Moigneu, C., Boillee, S., Couratier, P., Meininger, V., Pradat, P.F., Salachas, F., LeGuern, E., and Millecamps, S. (2014). Genetic analysis of SS18L1 in French amyotrophic lateral sclerosis. Neurobiol Aging 35.

Turner, M.R., Cagnin, A., Turkheimer, F.E., Miller, C.C.J., and Shaw, C.E. (2004). Evidence of widespread cerebral microglial activation in amyotrophic lateral sclerosis: an [C-11](R)-PK11195 positron emission tomography study. Neurobiol Dis 15, 601–609.

Van Deerlin, V.M., Leverenz, J.B., Bekris, L.M., Bird, T.D., Yuan, W., Elman, L.B., Clay, D., Wood, E.M., Chen-Plotkin, A.S., Martinez-Lage, M., et al. (2008). TARDBP mutations in amyotrophic lateral sclerosis with TDP-43 neuropathology: a genetic and histopathological analysis. The Lancet Neurology 7, 409–416.

Vance, C., Rogelj, B., Hortobagyi, T., De Vos, K.J., Nishimura, A.L., Sreedharan, J., Hu, X., Smith, B., Ruddy, D., Wright, P., et al. (2009). Mutations in FUS, an RNA processing protein, cause familial amyotrophic lateral sclerosis type 6. Science 323, 1208–1211.

Ward, C.L., Boggio, K.J., Johnson, B.N., Boyd, J.B., Douthwright, S., Shaffer, S.A., Landers, J.E., Glicksman, M.A., and Bosco, D.A. (2014). A loss of FUS/TLS function leads to impaired cellular proliferation. Cell Death Dis 5.

Weydt, P., Yuen, E.C., Ransom, B.R., and Moller, T. (2004). Increased cytotoxic potential of microglia from ALS-transgenic mice. Glia 48, 179–182.

Yamanaka, K., Boillee, S., Roberts, E.A., Garcia, M.L., McAlonis-Downes, M., Mikse, O.R., Cleveland, D.W., and Goldstein, L.S.B. (2008). Mutant SOD1 in cell types other than motor neurons and oligodendrocytes accelerates onset of disease in ALS mice. P Natl Acad Sci USA 105, 7594–7599.

Zhang, Y., Chen, K., Sloan, S.A., Bennett, M.L., Scholze, A.R., O’Keeffe, S., Phatnani, H.P., Guarnieri, P., Caneda, C., Ruderisch, N., et al. (2014). An RNA-sequencing transcriptome and splicing database of glia, neurons, and vascular cells of the cerebral cortex. J Neurosci 34, 11929–11947.

