## Supplementary Materials for "Loss of CREST leads to neuroinflammatory responses and ALS-like motor defects in mice"

**STAR METHODS**

**KEY RESOURCES TABLE**

| **REAGENT or RESOURCE** | **SOURCE** | **IDENTIFIER** |
| --- | --- | --- |
| **Antibodies** | | |
| CREST (Rabbit) | Willget | N/A |
| HA | Covance | Cat#MMS-101R, RRID: AB_291262 |
| β-Actin | Abmart | Cat#P30002M |
| BRG1 | Santa Cruz | Cat#sc-17796, RRID: AB_626762 |
| GAPDH | Abcam | Cat#ab8245, RRID: AB_2107448 |
| FUS | Abcam | Cat#ab124923, RRID: AB_10972861 |
| TDP43 | Cell Signaling | Cat#3448S, RRID: AB_2271509 |
| CREST (Goat) | Santa Cruz | Cat#sc-50912, RRID: AB_2195163 |
| Iba1 | Wako | Cat#019-19741, RRID: AB_839504 |
| ChAT | Millipore | Cat#AB144P, RRID: AB_2079751 |
| Neurofilament-L | Cell Signaling | Cat#2837, RRID: AB_823575 |
| Synapsin-1 | Cell Signaling | Cat#D1265 |
| YB1 | Abcam | Cat#ab76149, RRID: AB_2219276 |
| GFP | Invitrogen | Cat#A-11122, RRID: AB_221569 |
| SMI312 | BioLegend | Cat#837904, RRID: AB_2566782 |
| GFAP | Millipore | Cat#AB5541, RRID: AB_177521 |
| HDAC1 | Abcam | Cat#ab7028, RRID: AB_305705 |
| Donkey anti-rabbit HRP | GE-Healthcare | Cat#NA934V |
| Sheep anti-mouse HRP | GE-Healthcare | Cat#NA931V |
| Donkey anti-rabbit CF488A conjugate | Biotium | Cat#20015, RRID: AB_10559669 |
| Donkey anti-rabbit CF555 conjugate | Biotium | Cat#20038, RRID: AB_10558011 |
| Donkey anti-mouse CF555 conjugate | Biotium | Cat#20037, RRID: AB_10559035 |
| Donkey anti-goat CF555 conjugate | Biotium | Cat#20039, RRID: AB_10556967 |
| Goat anti-chicken CF555 conjugate | Biotium | Cat#20034, RRID: AB_10853135 |
| **Chemicals, Peptides, and Recombinant Proteins** | | |
| Papain | Worthington | Cat#LS003126 |
| Neurobasal medium | Gibco | Cat#21103-049 |
| B27 | Gibco | Cat#17504-044 |
| Protease inhibitor cocktail tablets | Roche | Cat#04693159001 |
| ECL Western Blotting Substrate | Pierce | Cat#32106 |
| Paraformaldehyde (PFA) | Sigma-Aldrich | Cat#P6148 |
| Triton X-100 | Sigma-Aldrich | Cat#T9284 |
| Trizol reagent | Invitrogen | Cat#15596-018 |
| Trichostatin A (TSA) | Cell Signaling | Cat#9950S |
| α-bungarotoxin (BTX) CF488A conjugate | Biotium | Cat#00005 |
| Sodium arsenite (SA) | Sigma-Aldrich | Cat#S7400 |
| **Critical Commercial Assays** | | |
| PrimeScript RT Master Mix | TaKaRa | Cat#RR036A |
| SYBR green premix | Toyobo | Cat#QPK-201 |
| KOD-Plus-Mutagenesis Kit | Toyobo | Cat#SMK-101 |
| ChIP Assay Kit | Upstate | Cat#17-295 |
| **Experimental Models: Organisms/Strains** | | |
| Mouse: C57BL/6 | JAX | RRID: IMSR_JAX: 000664 |
| **Oligonucleotides** | | |
| Forward primer for CREST Q388X cDNA mutagenesis:  5’-TAGGGCCAGTATGGAAATTACCAGCA-3’ | This paper | N/A |
| Reverse primer for CREST Q388X cDNA mutagenesis:  5’-TTCATAGCCGTAGGGCCGCTG-3’ | This paper | N/A |
| shRNA targeting sequence of mouse CREST:  5’-GGTCAGCAGTATGGAAGCT-3’ | Qiu and Ghosh, 2008 | N/A |
| shRNA targeting sequence of mouse BRG1:  5’-CCAAAGCAACCATCGAACT-3’ | Qiu and Ghosh, 2008 | N/A |
| CRISPR/Cas9 sgRNA sequence for KO mice:  5’-ATGCAGGGCCAGATCGGTAA-3’ | This paper | N/A |
| CRISPR/Cas9 sgRNA sequence for Q394X mice:  5’-GCGGCCTTACGGCTATGAAC-3’ | This paper | N/A |
| Repair donor for Q394X mice construction with CRISPR/Cas9:  5’-CATCGCAGACGGGACCTTCTGCCCAGCAGCAGCGGCCTTACGGCTATGAATGAGCAAGCTTTCTGGGCGTTTCAGGAAGCGCTATCTGCCAAGTGTCAAGTGA-3’ | This paper | N/A |
| Primer for qPCR: *Il1b* Forward:  5’-TGCCACCTTTTGACAGTGATG-3’ | This paper | N/A |
| Primer for qPCR: *Il1b* Reverse:  5’-TGATGTGCTGCTGCGAGATT-3’ | This paper | N/A |
| Primer for qPCR: *Tnfa* Forward:  5’-ACTTCGGGGTGATCGGTCCCC-3’ | This paper | N/A |
| Primer for qPCR: *Tnfa* Reverse:  5’-TGGTTTGCTACGACGTGGGCTAC-3’ | This paper | N/A |
| Primer for qPCR: *Cox-2* Forward:  5’-CCCTGCTGCCCGACACCTTC-3’ | This paper | N/A |
| Primer for qPCR: *Cox-2* Reverse:  5’-CCAGCAACCCGGCCAGCAAT-3’ | This paper | N/A |
| Primer for qPCR: *Cxcl10* Forward:  5’-AAGTGCTGCCGTCATTTTCT-3’ | This paper | N/A |
| Primer for qPCR: *Cxcl10* Reverse:  5’-CCTATGGCCCTCATTCTCAC-3’ | This paper | N/A |
| Primer for qPCR: *Ccl2* Forward:  5’-TTAAAAACCTGGATCGGAACCAA-3’ | This paper | N/A |
| Primer for qPCR: *Ccl2* Reverse:  5’-GCATTAGCTTCAGATTTACGGGT-3’ | This paper | N/A |
| Primer for qPCR: *Actb* Forward:  5’-GGCTCCTAGCACCATGAAGAT-3’ | This paper | N/A |
| Primer for qPCR: *Actb* Reverse:  5’-TAAAACGCAGCTCAGTAACAGT-3’ | This paper | N/A |
| QPCR Primer for ChIP: *Cxcl10* Forward:  5’-GCAATGCCCTCGGTTTACAG-3’ | This paper | N/A |
| QPCR Primer for ChIP: *Cxcl10* Reverse:  5’-TCTGCAAAGAGTTTCCCTCCC-3’ | This paper | N/A |
| QPCR Primer for ChIP: *Ccl2* Forward:  5’-CACTTCCTGGAAACACCCGA-3’ | This paper | N/A |
| QPCR Primer for ChIP: *Ccl2* Reverse:  5’-CTGCTCTGAGGCAGCCTTTT-3’ | This paper | N/A |
| QPCR Primer for ChIP: *c-Fos* Forward:  5’-GAAAGCCTGGGGCGTAGAG-3’ | This paper | N/A |
| QPCR Primer for ChIP: *c-Fos* Reverse:  5’-CCTCAGCTGGCGCCTTTAT-3’ | This paper | N/A |
| **Software and Algorithms** | | |
| ImageJ | NIH | RRID: SCR_003070 |
| GraphPad Prism 6 | GraphPad Software | RRID: SCR_002798 |
| EthoVision XT | Noldus | RRID: SCR_000441 |
| CatWalk XT | Noldus | N/A |
| StepOnePlus Real-Time PCR System | Applied Biosystems | RRID: SCR_015805 |
| DAVID Bioinformatics Resources | Leidos Biomedical Research, Inc. | RRID: SCR_001881 |
| **Databases** | | |
| Brain RNA-Seq | Zhang et al., 2014 | RRID: SCR_013736 |

**CONTACT FOR REANGENT AND RESOURCE SHARING**

**EXPERIMENTAL MODEL AND SUBJECT DETAILS**

The C57BL/6 *Mus musculus* was the experimental model in this study. All mice were maintained in a specific pathogen free (SPF) unit under constant temperature, humidity, ventilation and automatic light cycles. All genotypes, including CREST knockout and Q394X knock-in, used in experiments were confirmed by Sanger sequencing.

**METHOD DETAILS**

**Plasmid construction**

The human *CREST* coding sequence was synthesized and then subcloned to FUGW vector (Addgene Catalog #14883) with HA tag by enzyme digestion approach. We constructed the HA-Q388X mutant-expressing plasmid from HA-CREST WT-expressing plasmid with the KOD-Plus-Mutagenesis Kit (Toyobo) according to manufacturer’s instructions. The harboring vector of shRNA targeting mouse CREST or BRG1 is pFUGW-H1 empty vector (Addgene Catalog #25870). All the FUGW and pFUGW-H1 plasmids were packaged into lentivirus for high transfection efficiency in cultured primary neurons.

**Animals and ethics statement**

CREST knockout and Q394X knock-in mice were constructed with CRISPR/Cas9 system on the genetic background of C57BL/6 mouse. The sequences of small guide RNA (sgRNA) for CREST knockout or Q394X knock-in mice and the repair donor for Q394X knock-in mice are shown in the **KEY RESOURCES TABLE**. All animal-involved experiments were approved by the Biomedical Research Ethics Committee at the Shanghai Institutes for Biological Science (CAS). The use and care of animals were in accordance with the guidelines of this committee.

**Primary cortical neuron culture**

Cortices from embryonic day 15 to 16 C57BL/6 mice were digested with 20U/ml papain (Worthington, LS003126) at 37°C for 30 minutes. Cortical neurons were cultured within Neurobasal medium (Gibco, 21103-049) supplemented with 2% B27 (Gibco, 17504-044) at 37°C with proper density. We performed lentivirus transfection the day after plating and replaced the medium 24 hours after transfection. We collected neurons at 6 or 7 days *in vitro* for subsequent experiments.

**Western blotting**

The protein samples were harvested from cultured cortical neurons with RIPA buffer, and mouse brain cortices and lumbar spinal cords were homogenized in RIPA buffer (containing 150mM NaCl, 1% sodium deoxycholate, 0.1% SDS, 50mM Tris-HCl pH 7.4, 1% Triton X-100 and protease inhibitor cocktail tablets (Roche, 04693159001)). We sonicated the lysates with 10 sets of 30-second pulses on ice cold and the lysates were centrifuged at 13,000 rpm for 10 minutes at 4°C. The protein samples from the supernatant were run on 8-10% SDS-PAGEs at constant voltage and then transferred to PVDF membranes (Millipore, pore size: 0.45μm). Blots were blocked in 5% BSA in PBS-Tween for 2h at room temperature and then incubated with primary antibodies overnight at 4°C. After washing with PBS-Tween, the blots were incubated with secondary antibodies for 2h at room temperature. The protein bands were detected with chemiluminescence (ECL Western Blotting Substrate, Pierce, #32106).

**Immunohistochemistry**

For cultured primary neurons, we aspirated the medium and washed cells with PBS. 4% paraformaldehyde (PFA) in PBS was used to fix the cells for 20min at room temperature. After washing with PBS, cells were incubated in block buffer (3% BSA, 0.1% Triton X-100 in PBS) for 2h at room temperature. Then cells were incubated with primary antibodies overnight at 4°C, followed by the incubation of secondary antibodies for 2h at room temperature. For acquiring brain and spinal cord sections, animals were perfused transcardially with PBS then 4% PFA. After fixation in 4% PFA, brains and spinal cords were cut 40μm thick with cryostats sectioning of Leica CM1950. After washing in PBS, sections were incubated in block buffer (5% BSA, 0.3% Triton X-100 in PBS) for 2h at room temperature. Then sections were incubated with primary antibodies overnight at 4°C, followed by the incubation of secondary antibodies for 2h at room temperature. All images were captured on Nikon TiE-A1 plus confocal microscope.

**Morphological analysis**

We used Simple Neurite Tracer plugin in Image J software to quantify the total axon length of cultured neurons. For microglia morphology analysis, we used polygon selection tool to surround the projection of the cell bodies of Iba1-positive channel to calculate the soma area and roundness of microglia. Then we utilized the skeleton analysis method as previously reported (Morrison and Filosa, 2013) with minor modifications to estimate the length and number of branches per cell. In brief, we firstly visualized branches as many as possible, and then de-speckled noise signals to clear the background. After binary images were made, we used Skeletonize plugin in Image J, and then applied Analyze Skeleton plugin to calculate the length and number of skeletonized branches.

**RNA isolation and quantitative RT-PCR**

Total RNA was collected from neuron cultures or mouse tissue homogenates in Trizol reagent (Invitrogen, 15596-018) and extracted as manufacturer’s instructions. The reverse transcription was carried out with PrimeScript RT Master Mix (TaKaRa, RR036A), and 500-1000 ng total RNA was used per reaction. We performed real-time PCR with SYBR green premix (Toyobo, QPK-201) and analyzed data on the StepOnePlus Real-Time PCR System (Applied Biosystems). β-Actin was used as internal control.

**Chromatin immunoprecipitation**

ChIP Assay Kit (Upstate, 17-295) was used to perform ChIP assays according to the standard protocol in cultured cortical neurons. 200 base pairs ahead of the transcriptional start site were considered as the promoter region that interesting proteins bound. The sheared DNA samples were amplified and analyzed by real-time PCR. Signals were presented as the percentage of input. Anti-CREST (Willget) and anti-HDAC1 (Abcam, ab7028) antibodies were used in this experiment. IgG was the negative control.

**Behavior tests**

When CREST knockout mice and WT littermates were at the age of 12 months, rotarod tests were performed on a professional apparatus (Ugo Basile S.R.L.) every 2 weeks until 18 months. During this period, we used accelerating mode in which the rotating speed began at 8 rpm, and then elevated to 80 rpm in 300 seconds. At 18 months, we also performed fixed speed mode of rotarod tests on CREST knockout mice and WT littermates. In this mode, the rotating speed began at 20 rpm, and then lasted for 60 seconds. We also performed accelerating rotarod tests on Q394X knock-in mice at 12 or 18 months. We used two kinds of acceleration programs (4 to 40rpm, or 8 to 80 rpm in 300 seconds). We performed three trials in each session and measured the time mice could hold on the rotating rod to indicate the motor function. The apparatus setup of beam walking test was produced as previously described (Carter et al., 2001). We chose the square beam whose side length was 12 mm for training and test sessions. After three training days, we performed two consecutive trials on each mouse and recorded with video camera in the test session. We analyzed the beam traversing time and the slip times of hind paws of both CREST knockout and Q394X knock-in mice at 18 months.

In open field tests, mice were allowed to move freely in a 40cm × 40 cm white box for 10 minutes. We analyzed the video recordings by the Ethovision XT software (Noldus) to measure the distance mice moved and the time mice stayed in the center region. We used CatWalk XT system (Noldus) to analyze the gait and locomotion of knockout mice. We chose a region of 20 cm length as the capture zone, and footprints could be visualized and analyzed when mice traversed this zone. The grip strength apparatus (Ugo Basile) was applied to measure the forelimb strength of knockout mice at 18 months.

**QUANTIFICATION AND STATISTICAL ANALYSIS**

We used Graph Pad Prism 6 software (La Jolla) as the statistics tool to determine whether there were statistically significant differences between groups. Student’s t test was used in the analysis between two groups. One-way ANOVA was used between three groups. Two-way ANOVA was used in the analysis consisted of two factors. The meanings of p values are indicated as blow: *p<0.05, **p< 0.01, ***p< 0.001, and ****p<0.0001.
